# The mechanistic basis for chromatin invasion and remodeling by the yeast pioneer transcription factor Rap1

**DOI:** 10.1101/541284

**Authors:** Maxime Mivelaz, Anne-Marinette Cao, Slawomir Kubik, Sevil Zencir, Ruud Hovius, Iuliia Boichenko, Anna Maria Stachowicz, Christoph F. Kurat, David Shore, Beat Fierz

## Abstract

Pioneer transcription factors (pTFs) bind to target sites within compact chromatin initiating chromatin remodeling and controlling the recruitment of downstream factors. The mechanisms by which pTFs overcome the chromatin barrier are not well understood. Here we reveal, using single-molecule fluorescence approaches, how the yeast transcription factor Rap1 invades and remodels chromatin. Using a reconstituted chromatin system replicating yeast promoter architecture we demonstrate that Rap1 can bind nucleosomal DNA within a chromatin fiber, but with shortened dwell times compared to naked DNA. Moreover, we show that Rap1 binding opens chromatin fiber structure by inhibiting nucleosome-nucleosome contacts. Finally, we reveal that Rap1 collaborates with the chromatin remodeler RSC to destabilize promoter nucleosomes, paving the way to form long-lived bound states on now exposed DNA. Together, our results provide a mechanistic view of how Rap1 gains access and opens chromatin, thereby establishing an active promoter architecture and controlling gene expression.

---

Chromatin acts as a barrier for proteins which require access to the DNA. Both the target search and binding-site recognition of transcription factors (TFs) are restricted by the presence of nucleosomes and chromatin higher-order structure^1, 2^, resulting in reduced binding rates and residence times compared to free DNA^3^. However, a subset of transcription factors, named ‘pioneer transcription factors’ (pTFs)^4^, have the ability to invade compact chromatin domains, where they can find and bind to their cognate recognition motifs. They then initiate an opening of chromatin structure^5, 6^, which can coincide with linker histone loss^7^ or removal of certain labile nucleosomes^8-10^. Such remodeled chromatin is accessible to subsequent non-pioneer TFs^5^, which enact changes in transcriptional programs^11, 12^.

A common feature of DNA binding domains (DBDs) found in pTFs is their ability to bind partial sequence motifs displayed on the surface of a nucleosome^12^. The presence of nucleosomes may therefore have limited effects on both on-rates and residence times of pTFs. Beyond the nucleosome, higher-order chromatin structure further constrains DNA conformation and accessibility^13^. High-resolution structural studies on reconstituted chromatin revealed that local structural elements such as tetranucleosome units form the basis of chromatin fiber organization^14^. Genomic studies have confirmed the prevalence of tetranucleosome contacts *in vivo*, employing Micro-C^15^ or *in situ* radical fragmentation of chromatin^16^. Neighboring tetranucleosome units can interact and form fiber segments with two intertwined stacks of nucleosomes^14, 17, 18^. Such local structure may therefore restrict DNA access with unknown consequences for pTF interaction dynamics.

It is not well understood how pTFs probe DNA sequences within chromatin as well as how they invade and subsequently remodel chromatin structure. The intrinsic dynamics of chromatin itself however might provide a potential mechanism for pTF invasion^19^. Recent studies using force spectroscopy^17^ or single-molecule FRET^20^ to probe chromatin fiber dynamics revealed conformational exchange processes acting over multiple time scales, from microseconds to seconds. It is thus conceivable that pTFs exploit dynamic site exposure within chromatin fibers to invade compact chromatin, where they then recruit additional cellular machinery to enact necessary conformational reorganization to alter gene expression (**Fig. 1a**). Currently, direct experimental insight into such mechanisms is lacking.

**Figure 1.**
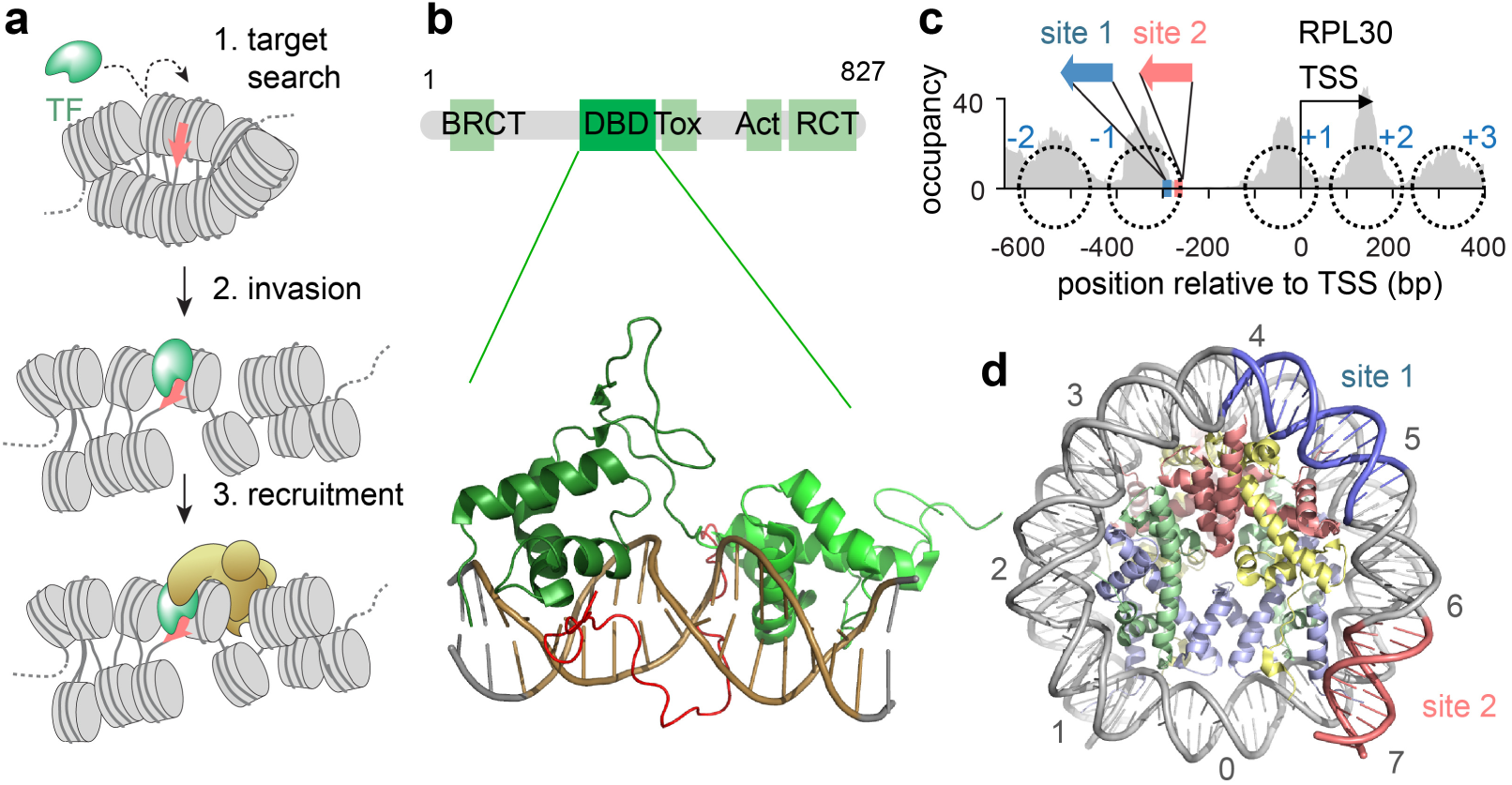
Rap1 as a pioneer factor in budding yeast. **a)** Scheme of pTF function: After a target search (1) for its cognate binding site, the pTF invades compact chromatin structure (2), followed by local chromatin opening and recruitment of the transcription machinery (3). **b)** Domain organization of budding yeast Rap1 (above) and X-ray crystal structure of Rap1 in complex with its cognate DNA motif (PDB code 3ukg, ref. ^68^). Rap1 is constituted of several regions including a BRCA 1 C Terminus (BRCT), DNA Binding Domain (DBD), Toxicity region (Tox), Transcription Activation domain (Act) and the Rap1 C-Terminus (RCT). **c)** The organization of the *RPL30* promoter. Grey: MNase-seq profile (Rap1 depleted by anchor-away)^25^ reveals nucleosome positions in the absence of Rap1 (black dotted circles). Plotted is nucleosome occupancy reads, normalized to 10^7^ total reads. The Rap1 binding site 1 (*S1*, high affinity) and site 2 (*S2*, medium affinity) fall on the −1 nucleosome. **d)** Promoter −1 nucleosome, showing Rap1 binding sites *S1* and *S2* mapped on the DNA (PDB code 1AOI, ref. ^69^). The numbers indicate super helical locations (SHL) of the nucleosomal DNA.

Here, we reveal the mechanism of chromatin invasion, target binding and chromatin remodeling of the pTF Rap1 (repressor <u>a</u>ctivator <u>p</u>rotein 1). Rap1 is an important general regulatory factor (GRF) of transcription in budding yeast^10^. It plays multiple roles *in vivo* including the transcriptional regulation of around 5% of yeast genes^21^ and the maintenance of telomeric integrity^22^. The Rap1 DNA binding domain (DBD) consists of dual Myb-type domains which are connected by a short unstructured linker^23^ (**Fig. 1b**). The DBD binds a class of 13 bp consensus motifs with high affinity (**Supplementary Fig. 1a**), only requiring direct access to one face of the DNA (**Fig. 1b**). Importantly, Rap1 has been shown to be able to engage a single motif in multiple binding modes, involving one or both Myb-domains^24^, thus giving the pTF flexibility in DNA recognition.

A particularly important gene family co-regulated by Rap1 are ribosomal protein genes. Rap1 binds to the promoter/enhancer regions of >90% of these genes and it initiates the recruitment of combinations of additional TFs, including Hmo1, Fhl1 and Ifh1^10^. In one of the two largest categories of ribosomal protein genes (category I), two closely spaced Rap1 binding sites are situated upstream of the transcription start site (TSS)^10^. When Rap1 is depleted from the nucleus and its target genes are down-regulated, the Rap1 binding sites are covered by a stable nucleosome^25^. In the presence of Rap1, a large nucleosome-depleted region (NDR) forms^10, 25^. Such NDRs are typical for most active eukaryotic promoters^26^, and depend on the action of remodeling factors, including RSC^27-33^, SWI/SNF^34, 35^ and INO80^36^.

Digestion of yeast chromatin with limited amounts of micrococcal nuclease (MNase) followed by sequencing (MNase-seq)^37^ revealed that many NDRs contain MNase-sensitive particles^38-41^, which comprise protein complexes^42^ and may correspond to destabilized promoter nucleosomes^25, 32, 33^. In category I promoters such MNase sensitive nucleosome-like particles appear upstream of the +1 nucleosome, co-existing with bound Rap1^25^. Thus, Rap1 binds target sites within the promoter nucleosome, which is subsequently destabilized, allowing the recruitment of further TFs followed by the transcription machinery. The mechanism by which Rap1 finds its target in compacted chromatin and how it subsequently acts to open chromatin and to destabilize promoter nucleosomes are however not understood.

In this study we dissect the mechanism of Rap1 invasion into compact chromatin. We reconstituted nucleosomes and chromatin fibers, containing Rap1 binding sites in the configuration found in category I promoters. We find that the residence times, but not binding rates, of Rap1 are strongly reduced by the presence of both nucleosomes and, to an increased extent, chromatin fiber structure. Importantly, we show that Rap1 binding does not disrupt or decidedly alter nucleosome structure. In contrast, our single-molecule FRET measurements directly reveal that invasion of the chromatin fiber by Rap1 results in local opening of chromatin structure. Finally, we demonstrate that Rap1 binding tags nucleosomes for eviction by RSC. The remodeled chromatin structure and the destabilized nucleosomes then provide an opening for stable Rap1 binding, subsequent factor access and finally ribosomal gene activation.

## RESULTS

### Rap1 binds to nucleosomes via non-specific and specific DNA interactions

In a first set of experiments we investigated if Rap1 can bind target sequences in nucleosomal DNA and, if this is the case, through which mechanism. A typical example of a Rap1 controlled category I ribosome protein promoter is the ribosomal protein L30 (*RPL30*) promoter (**Fig. 1c**). Using MNase-seq under Rap1-depleted conditions, we precisely mapped the position of the −1 nucleosome at this promoter (**Fig. 1c**), which contains two Rap1 binding sites and which is destabilized *in vivo* upon Rap1 binding^25^. This allowed us to establish the relative orientation of the two Rap1 target sites on the promoter nucleosome structure: site 1 (*S1*) is located on the nucleosome near super helical location (SHL) 4.5, whereas site 2 (*S2*) resides near the DNA entry-exit site at SHL 6.5 (**Fig. 1d**). Importantly, on naked DNA Rap1 shows different affinity for the two sites. Electrophoretic mobility shift assays (EMSA) revealed that Rap1 binds to site 1 with an affinity of ∼10 nM, whereas binding to site 2 is of weaker affinity (∼30 nM) (**Supplementary Fig. 1c-e**). *In vivo,* both sites contribute to the expression of *RPL30*^10^. To directly observe dynamic Rap1 binding to promoter nucleosomes, we used a single-molecule total internal reflection fluorescence microscopy approach (smTIRFM) that reveals TF binding to immobilized nucleosomes or DNA via fluorescence colocalization (**Fig. 2a**)^43^. Several reagents were required: First, we generated a 200 base pair (bp) 601 DNA template^44^ which contained one of the two Rap1 binding sites, *S1* or *S2,* positioned so as to match their location on the *RPL30* promoter when reconstituted into nucleosomes (**Fig. 1d**, **Supplementary Fig. 1b** and **Supplementary Tables 1-3**). Moreover, the DNA constructs contained a fluorescent dye (Alexa Fluor 647) for detection in the far-red channel and a biotin moiety for immobilization. We then either used this DNA directly or reconstituted nucleosomes using recombinantly expressed histones (**Fig. 2a** and **Supplementary Fig. 2**). Second, we purified full-length Rap1 as a Halo-tag fusion from insect cells, using a baculovirus expression system and maltose binding protein (MBP) as a solubility tag. After expression and MBP removal, Rap1 was fluorescently labeled with the highly photostable dye JF-549^45^ (allowing detection in the green-orange channel) and purified by size exclusion chromatography (**Fig. 2b**, Supplemental Fig. 3). Importantly, labeled Rap1 exhibited similar DNA binding compared to published values^10^ (**Supplementary Fig. 1c-e**).

**Figure 2.**
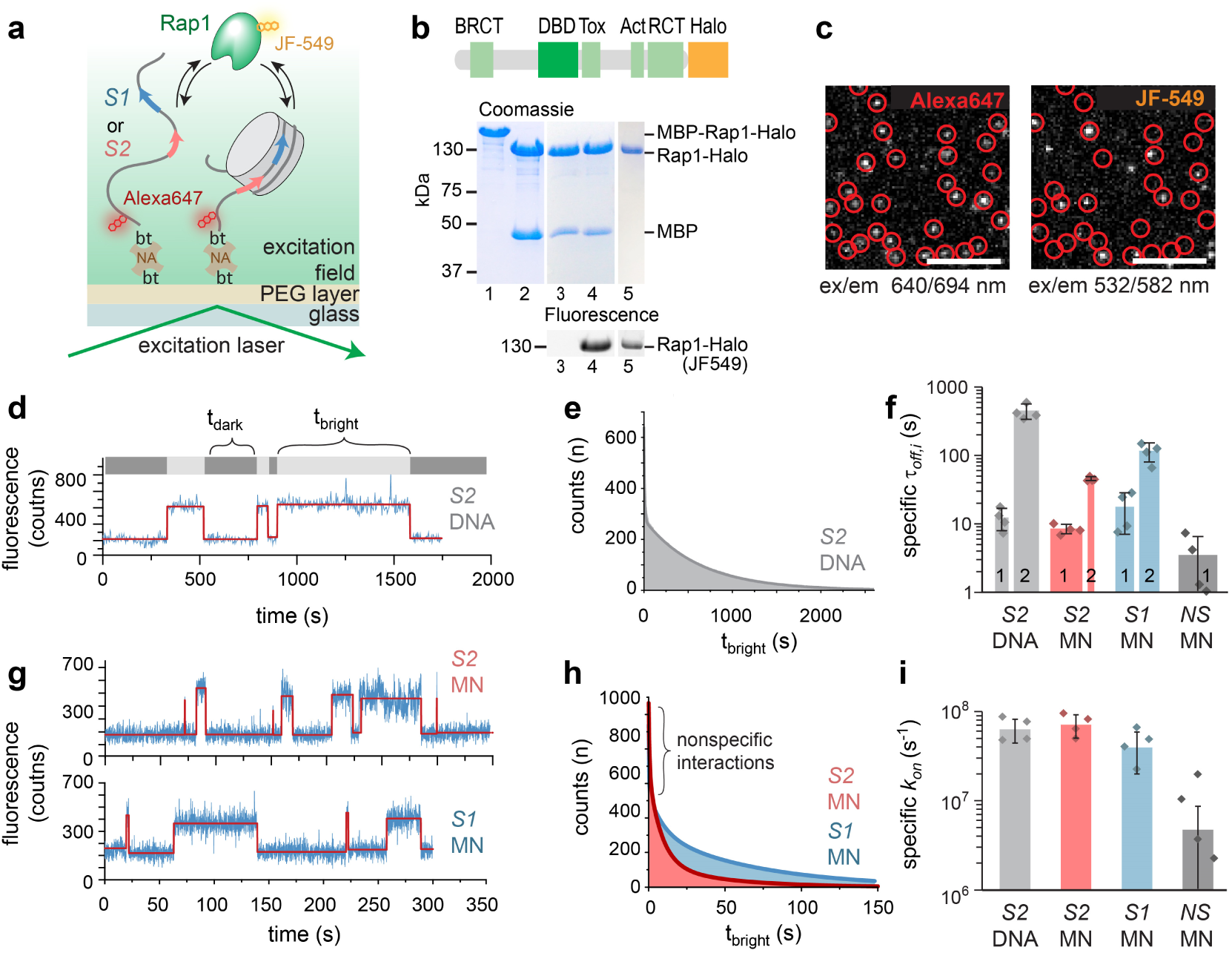
Rap1 recognizes target sites within nucleosomal DNA. **a)** Schematic view of the smTIRFM experiment, probing Rap1 binding to immobilized DNA or nucleosomes (using biotin-neutravidin (bt-NA) chemistry), containing either *S1* (high affinity) or *S2* (medium affinity) and labeled with Alexa Fluor647. **b)** Expression of JF-549 labeled Rap1, using Halo-tag. Lanes: 1. Purified MBP-Rap1-Halo construct; 2. MBP cleaved; 3. Before JF-549 labeling; 4. After JF-549 labeling; 5. Purified, labeled Rap1 construct. **c)** Representative smTIRF images showing nucleosome positions in the far red channel (left, red-circles) and Rap1 interaction dynamics in the green-orange channel (right). Scale bar: 5 μm; ex, excitation wavelength; em, emission wavelength. **d**) Representative fluorescence time trace of Rap1 binding events to *S2* containing free DNA, detected by JF-549 emission. The trace is fitted by step function (red) and *t_dark_* and *t_bright_* were determined by a thresholding algorithm. **e**) Cumulative histogram of Rap1 binding intervals (*t_bright_*) on *S2* DNA fitted by a 2-exponential function 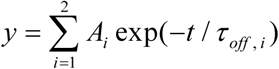 (solid line). For fit results, see **Table 1**. **f**) Specific dissociation time constants (*r*_off,i_ > 1 s) of Rap1 for *S2* DNA, *S1* and *S2* containing mono-nucleosomes (MN) or nucleosomes lacking a binding site (601), uncorrected for dye photobleaching. The width of the bars indicate the percentage of events associated with the indicated time constants (i.e. amplitudes *A_i_*of the multi-exponential fits shown in **e**,**h**). N = 4-5, error bars: s.d. **g**) Representative fluorescence time trace of Rap1 binding events to *S1* (bottom) and *S2* (top) containing MNs. The data was analyzed as in **d**. **h**) Cumulative histogram of Rap1 binding intervals (*t_bright_*) on *S1* and *S2* containing MNs fitted by a 3-exponential function 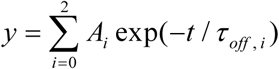 (solid line). For fit results, see **Table 1** and panel **f**). **i**) Specific on-rates (*k_on_ = 1/r_on_*), for all species obtained from a single-exponential fit to cumulative histograms of *t_dark_* values, and corrected for the contribution from non-specific interactions (**Materials & Methods**).

Having all components in hand, in a first set of experiments we immobilized *S1-* or *S2*-containing naked DNA in a microfluidic channel under native ionic strength (130 mM KCl) and determined their position by smTIRFM imaging in the far-red channel (**Fig. 2c**). We then injected JF-549-labeled Rap1 molecules at a concentration chosen such that individual, non-overlapping binding events could be detected as fluorescent spots in the green-orange channel (usually 50-100 pM). Target binding was observed when Rap1 detections colocalized with DNA positions (**Fig. 2c**). We then recorded movies (using a frame rate of 0.6 s), which revealed binding kinetics of Rap1 to *S1-* or *S2*-containing naked DNA. For each DNA localization, extracted kinetic traces allowed us to determine directly the length of individual binding events (*t_bright_*) and intermittent search times (*t_dark_*). We further limited the effect of dye photobleaching on residence time measurements by stroboscopic imaging (**Supplementary Fig. 4a** and Table 1).

**Table 1:**
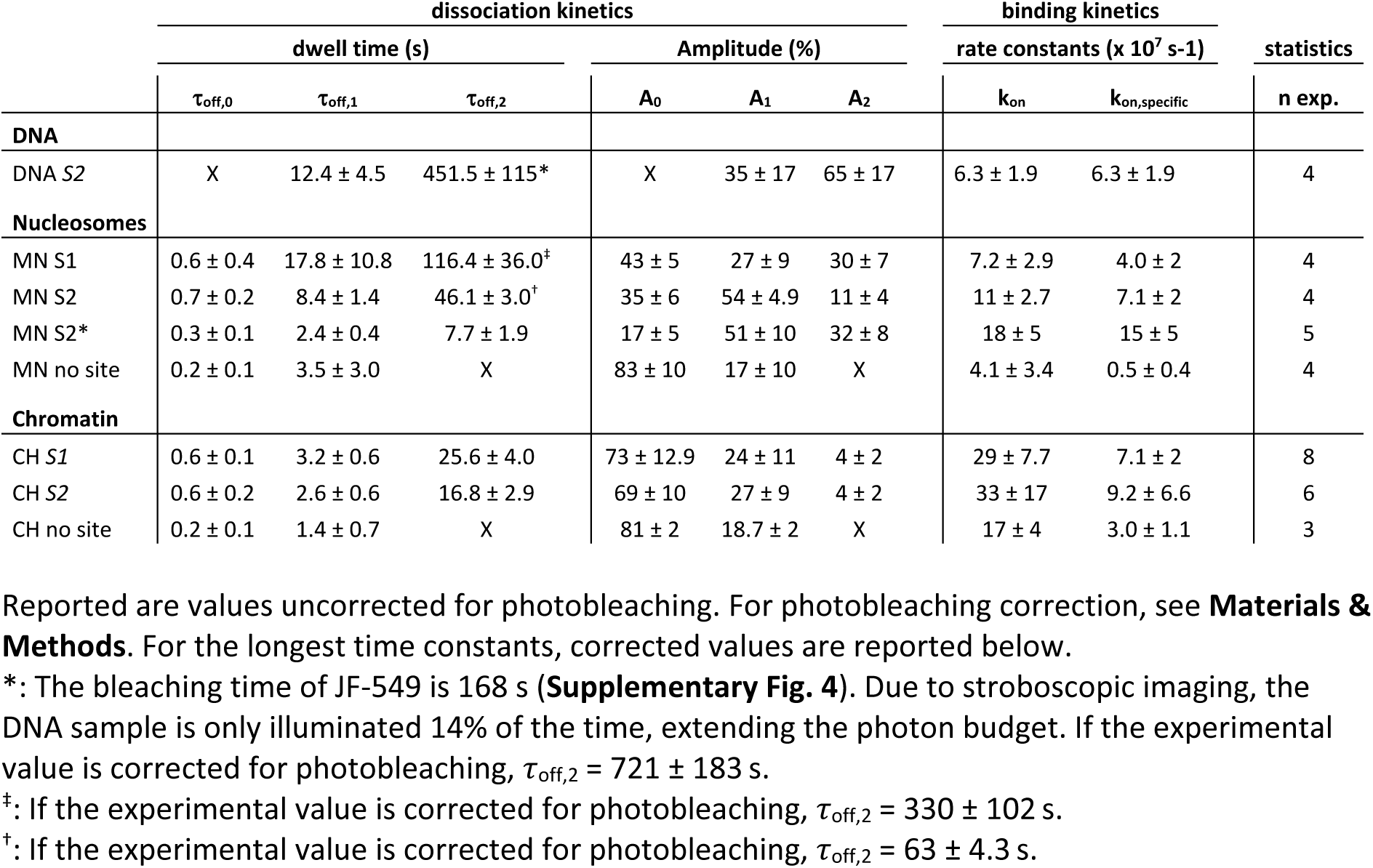
All kinetic parameters of Rap1 interacting with DNA, nucleosomes and chromatin.

While dynamic Rap1 binding was observed for the medium affinity site *S2* (**Fig. 2d**), individual binding events to the high affinity site *S1* were so long (> 40 min) that we were not able to obtain suitable statistics (**Supplementary Fig. 4b**). For *S2*-containing DNA, we then constructed cumulative lifetime histograms of bright times (*t_bright_*) (**Fig. 2e**), which were fitted using a bi-exponential function, yielding two residence times τ_off,1_ and τ_off,2_ (**Fig. 2f** and Table 1). Of all binding events, 35% exhibited a short residence time (τ_off,1_ = 12.4 ± 4.5 s) whereas the remaining 65% showed slow Rap1 dissociation kinetics (τ_off,2_ = 452 ± 115 s). Due to the dual Myb-type DBD, these different residence times may indicate different binding modes where either the entire or only a partial DNA binding motif is engaged. Under equilibrium binding conditions, Rap1 thus forms highly stable complexes with free DNA, resulting in residence times in the minutes to hours range for *S1* and *S2*.

In contrast, the presence of nucleosomes critically shortened the residence time of Rap1, as clearly observed in kinetic traces (recorded with a framerate of 10 Hz) for mononucleosomes (MNs) containing either *S1* or *S2* (**Fig. 2g**), and in the corresponding lifetime histograms (**Fig. 2h**). Here, a tri-exponential function was required to describe the data (**Supplementary Fig. 5**). Around 50% of all detected events were short-lived, with a time constant of 0.2 < τ_off,0_ < 0.7 s (Table 1). We attribute these fast events to non-specific probing interactions of nucleosomal DNA. Rap1 binding to *S1* or *S2* further revealed two additional longer time constants τ_off,1_ and τ_off,2_: Rap1 binding to the high affinity site *S1* was associated with longer residence times (τ_off,1_ = 18 ± 11 s and τ_off,2_ > 100 s) compared to *S2* (τ_off,1_ = 8.4 ± 1.4 s, τ_off,2_ = 46 ± 3 s) (**Fig. 2f**). This is not necessarily expected, as *S1* is located further within the nucleosome structure and thus potentially less accessible than *S2*, which resides at the DNA entry-exit site. To test the effect of site positioning on the nucleosome, we thus moved *S2* from SHL 6.5 to SHL 4.5, further within the nucleosome structure. Indeed, moving the site resulted in an additional reduction in Rap1 residence time to τ_off,1_ = 2.4 ± 0.4 s and τ_off,2_ = 7.7 ± 1.9 s (**Supplementary Fig. 4c-e**). Of note, under our measurement conditions (yielding individual, non-overlapping Rap1 binding events) having both binding sites *S1* and *S2* within the same nucleosomes resulted in a superposition of the individual binding kinetics (**Supplementary Fig. 4l,m**). Higher Rap1 concentrations or combinatorial binding with additional TFs might result in cooperative effects, which will require further investigations in the future.

Specific binding rates (*k_on_*), which were obtained from analyzing lifetime histograms of dark times (*t_dark_*) (**Supplemental Fig. 4f-h**, for the calculation of specific binding rates see **Materials & Methods**), were comparable for all DNA and nucleosome constructs analyzed (**Fig. 2i**). This demonstrates that the Rap1 target search kinetics are influenced by the presence of nucleosomes or the positioning of the tested binding sites. Finally, we also probed Rap1 binding to nucleosomes which did not contain any binding sites (**Supplementary Fig. 4i,k**). The large majority (83%) of all binding events detected were shorter than a second, while the remaining 17% persisted for only 3.5 ± 3 s (**Fig. 2f,i**).

Together, these results indicate that Rap1 can bind to nucleosomal DNA, with overall similar on-rates and with strongly (> 10-fold) reduced residence times compared to free DNA, which depend on the presence of a binding motif and its position on the nucleosome.

### Chromatin structure shortens Rap1 dwell times

pTFs not only need to find their target sites within nucleosomal DNA, but they have to invade compact chromatin structure. Indeed, compact chromatin, and in particular heterochromatin, has been shown to reduce pTF accessibility^46^. We therefore proceeded to investigate the mechanism of dynamic chromatin structure invasion by Rap1. To this end, we employed a modular system to construct chromatin fibers^20^, based on a 12-mer repeat of 601 nucleosome positioning sequence each separated by 30 bp linker DNA. Our modular approach involves a series of preparative DNA ligations followed by purification steps and allows us to include modified DNA sequences or fluorescent probes at distinct sites^20^. We assembled two chromatin fiber types, containing Rap1 target sites *S1* or *S2* in their central nucleosome (N6) at the same location as in the *RPL30* −1 nucleosome (**Fig. 3a**, **Supplementary Fig. 6-7**). The chromatin fibers (which further contained the dye Alexa647 for localization) were then immobilized in a flow cell and Rap1 binding dynamics were determined using our smTIRFM approach under native salt conditions (130 mM KCl, **Fig. 3b**). Importantly, under these ionic strength conditions chromatin fibers exist in a compact state^47^ (see also below). Compared to mononucleosomes, the observed dynamic traces for Rap1 binding to both *S1-* and *S2*-containing chromatin fibers exhibited an increase in short (0.6 s) binding events (∼70% of all detections, **Fig. 3c**), which can be attributed to non-specific Rap1-DNA probing interactions. Rap1 thus rapidly samples the chromatin fiber in its search for potential target sites.

**Figure 3.**
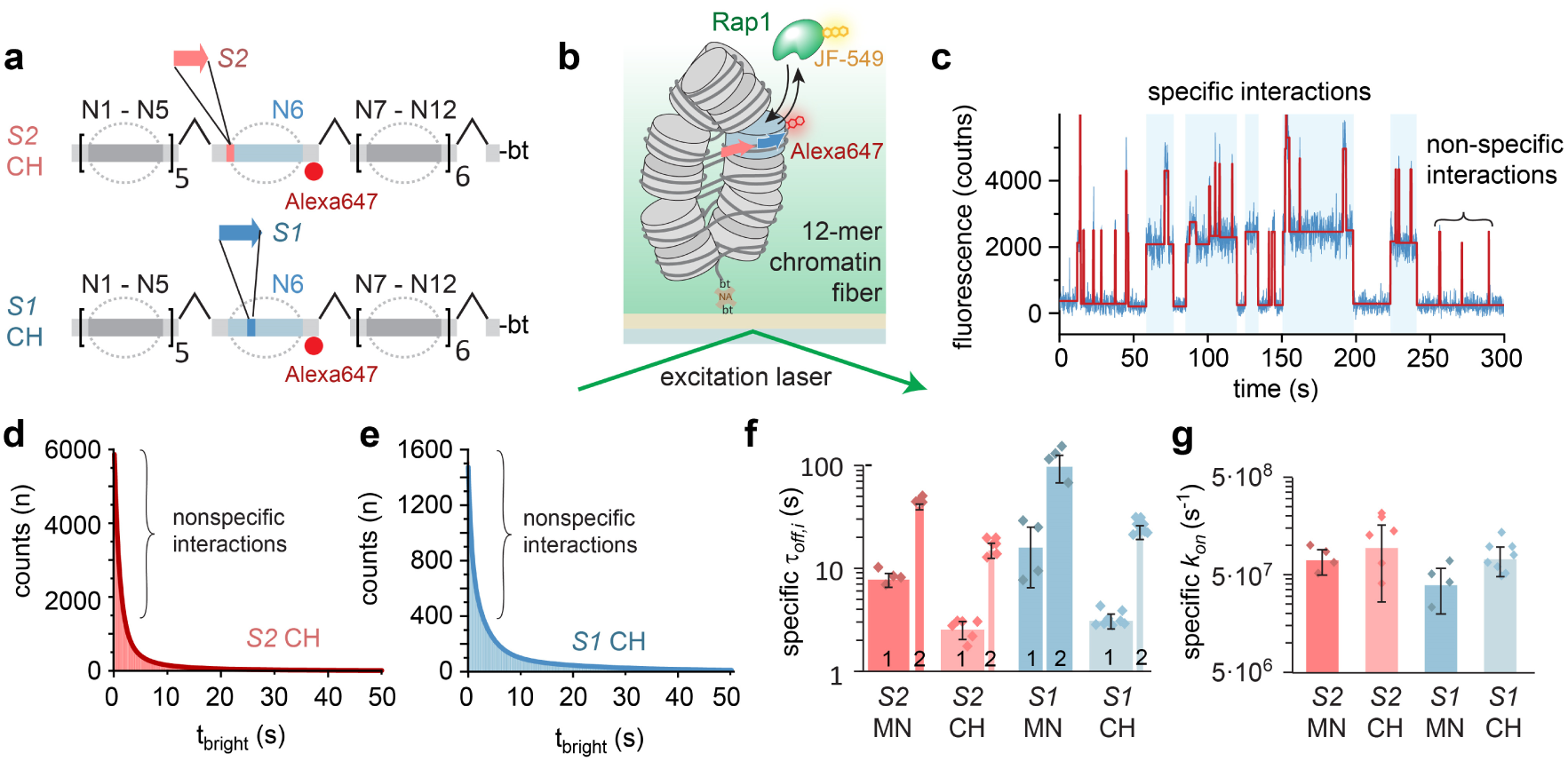
Chromatin higher-order structure reduces Rap1 dwell time. **a**) Schematic view of the preparative DNA ligation used to introduce Rap1 target sites *S1* and *S2* into the central nucleosomes (N6) in a chromatin fiber (CH). **b**) Scheme of the smTIRFM experiment to measure Rap1 binding kinetics in a chromatin fiber context. **c**) Representative fluorescence time trace of Rap1 binding events to *S1* containing chromatin arrays. The trace is fitted by step function (red) and t_dark_ and t_bright_ were determined by a thresholding algorithm. **d**) Cumulative histogram of Rap1 binding intervals (t_bright_) to chromatin fibers, containing *S1* fitted by a 3-exponential function (solid line). For fit results, see **Table 1**. **e**) Cumulative histogram of Rap1 binding to chromatin arrays, containing *S2* fitted by a 3-exponential function (solid line). For fit results, see **Table 1**. **f**) Specific binding time constants (*r*_off,i_ > 1 s) of Rap1 for S1 in a nucleosome (MN) vs. chromatin fiber (CH) and S2 MN vs CH. The width of the bars indicate the percentage of events associated with the indicated time constants (i.e. amplitudes *A_i_* of the multi-exponential fits shown in **d**,**e**). N = 4-5, error bars: s.d. **g**) Specific on-rates (*k_on_ = 1/r_on_*), for mononucleosomes (MN) and chromatin fibers (CH) containing *S1* and *S2*, obtained from a single-exponential fit to cumulative histograms of *t_dark_* values, and corrected for the contribution from non-specific interactions (**Materials & Methods**).

At a lower frequency, long-lived specific binding events were detected (**Fig. 3c**), demonstrating that Rap1 can indeed invade compact chromatin fiber and bind to its target site. In contrast, in chromatin fibers without binding sites mostly short-lived nonspecific binding events were observed (**Supplementary Fig. 8** and **Table 1**). Analyzing lifetime histograms for Rap1 binding to chromatin fibers containing either *S2* (**Fig. 3d**) or *S1* (**Fig. 3e**) revealed that the longer-lived specific binding events are governed by complex kinetics, exhibiting two time constants (**Table 1**). This is similar to the situation in mononucleosomes and most likely reflects multiple Rap1 binding modes. Intriguingly, compared to nucleosomes the residence times of Rap1 were substantially reduced in chromatin fibers (**Fig. 3f**). For *S2*, both time constants were reduced about 3-fold (τ_off,1_ = 2.6 ± 0.6 s, and τ_off,2_ = 16.8 ± 3 s), whereas for the high-affinity site *S1* a 5-fold reduction in dwell times was observed (τ_off,1_ = 3.2 ± 0.6 s, and τ_off,2_ = 25.6 ± 4.0 s). This shortening of Rap1 dwell times demonstrates that chromatin fiber structure acts as an additional barrier to Rap1 binding.

To determine if chromatin also inhibits the target search process of Rap1 we investigated the on-rates (*k_on_*) in chromatin compared to nucleosomes. Surprisingly, we could not detect any significant reduction in the Rap1 binding rate for chromatin fibers containing *S1* or *S2* (**Fig. 3g**). It is thus conceivable that Rap1 can hop or slide along chromatin in search of the appropriate binding site, using non-specific DNA interactions as a means of chromatin anchoring. Chromatin dynamics on the ms time-scale^20^ will eventually expose internal DNA sites, allowing the factor to bind to its target sequence.

### Rap1 binding does not evict or distort bound nucleosomes

Having established that Rap1 indeed binds to nucleosomes and can invade chromatin structure (albeit with diminished affinity), we wondered if the pTF can remodel chromatin, i.e. by directly opening chromatin structure^11^. In cells, Rap1 binding results in the destabilization and disruption of promoter nucleosomes^10, 25^, thereby paving the way for binding of subsequent TFs and establishing a chromatin state permissive to transcription. We wondered if Rap1 association actually destabilizes nucleosome structure, and leads to DNA unwrapping as observed for other TFs^48^. We therefore established a FRET-based assay to monitor nucleosomal DNA unwrapping as a function of Rap1 binding (**Fig. 4a**, **Supplementary Fig. 9**). We positioned FRET donor (Alexa568) and acceptor (Alexa647) dyes within the linker DNA of *S1-* and *S2*-containing nucleosomes, such that partial DNA unwrapping (or nucleosome disassembly) will lead to FRET loss (**Fig. 4a,b**). When titrating Rap1, complexes were formed containing several Rap1 molecule per nucleosomes at the highest concentrations, as judged by EMSA (**Fig. 4c**). No change in FRET efficiency (*E_FRET_*) was observed for either *S1* or *S2* nucleosomes (**Fig. 4d,e**) even at the highest Rap1 concentrations (**Fig. 4f**). Importantly, for TFs that envelop their target sequence, e.g. LexA or Gal4, nucleosome unwrapping was observed in similar, 601-based nucleosome systems^3, 48, 49^. These experiments directly show that Rap1 binding to *S1* or *S2* does not affect nucleosome structure and does not result in DNA unwrapping or histone loss.

**Figure 4.**
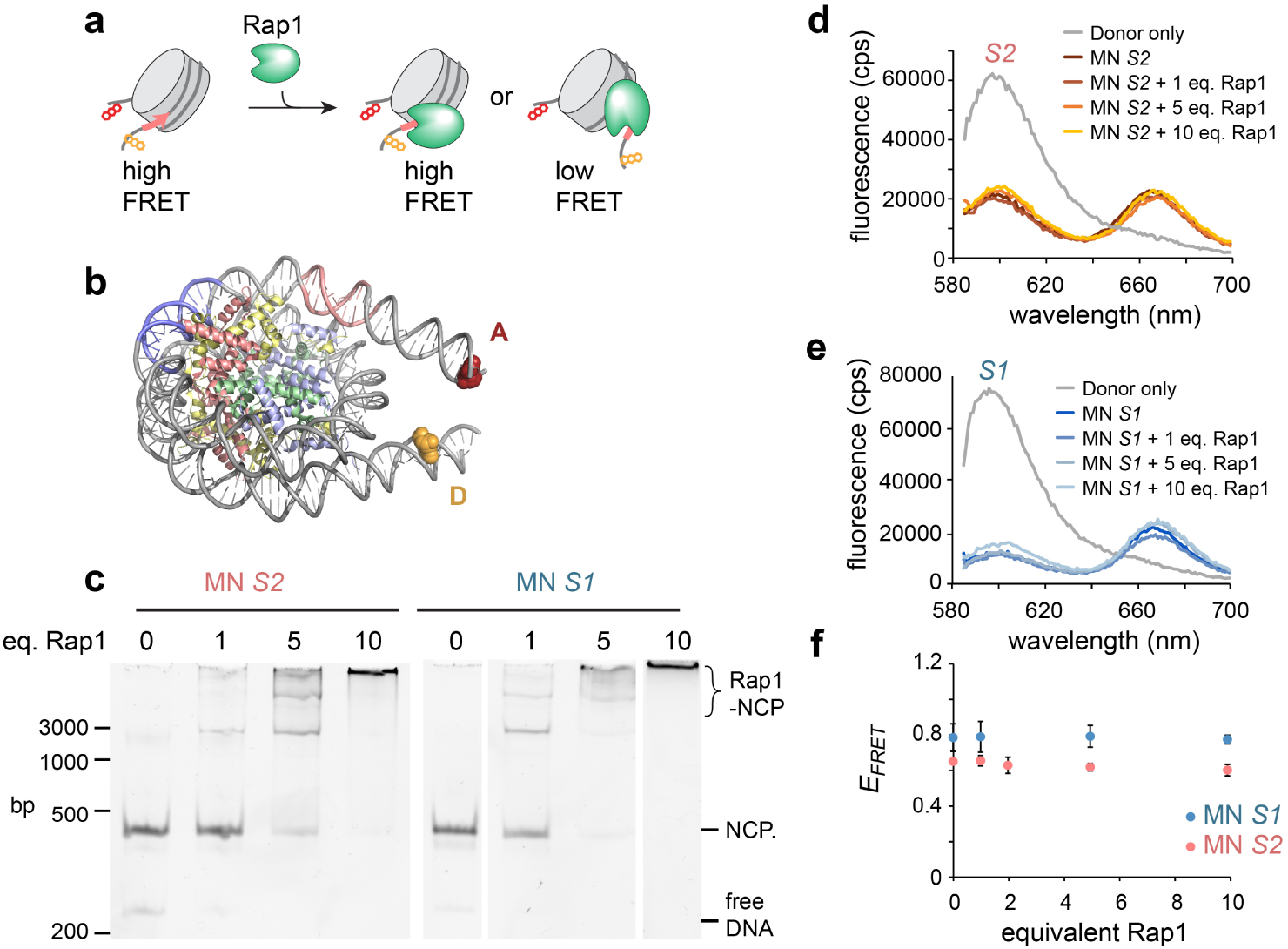
Rap1 does not open nucleosome structure. **a**) Scheme of a FRET approach to probe nucleosome structure as a function of bound Rap1. **b**) Nucleosome structure (PDB code 1AOI) showing attachment points of FRET probes. **c**) EMSA showing Rap1 binding to *S1* and *S2* nucleosomes at indicated concentration equivalents (eq.). **d**) Fluorescence spectra for *S2* nucleosome in complex with indicated equivalents of Rap1. **e**) Fluorescence spectra for *S1* nucleosome in complex with indicated equivalents of Rap1. **f**) FRET efficiency calculated for *S2* and *S1* nucleosomes as a function of equivalents added Rap1. Error bars: s.d., n = 2.

### Rap1 locally opens chromatin structure

While the structure of individual nucleosomes is not disrupted by Rap1 binding, higher-order chromatin structure might be altered. To directly determine the effect of Rap1 binding on chromatin structure, we used a FRET scheme that directly reports on nucleosome stacking interactions^20, 50^. To this end, we inserted a Rap1 binding site (*S2*) in the central nucleosome (N6) in a 12-mer chromatin fiber. Using our modular chromatin assembly system^16^, we further flanked this central nucleosome by nucleosomes carrying a FRET donor (Cy3B in N5) and FRET acceptor (Alexa647 in N7) (**Fig. 5a**, **Supplementary Fig. 10-11**). Chromatin compaction decreases the distance between donor and acceptor dyes (**Fig. 5b**), resulting in an increase in FRET. Conversely, a local loss in higher-order structure (e.g. due to unstacking of a tetranucleosome unit) will result in a distinct reduction in *E_FRET_*.

**Figure 5.**
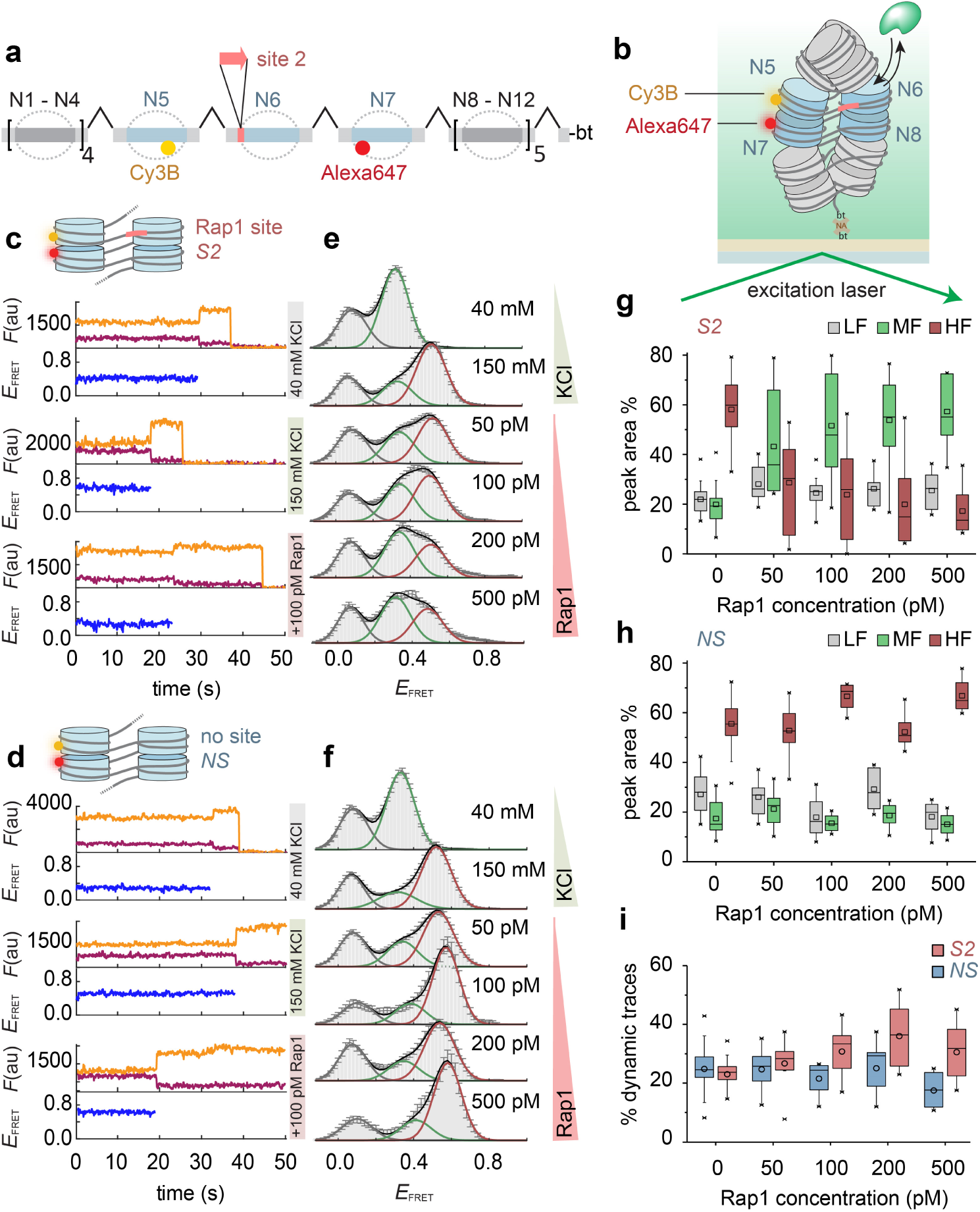
Chromatin remodeling induced by Rap1 invasion as observed by smFRET. **a**) Scheme of the assembly of 12-nucleosome chromatin fibers containing a Rap1 site at nucleosome N6, as well as a FRET donor (Cy3B, yellow) and acceptor (Alexa647, red) at nucleosomes N5 and N7. **b**) Scheme of a smFRET-TIRF experiment allowing to monitor the interaction between Rap1 and surface-immobilized single chromatin fibers. **c**) Individual kinetic traces of donor (orange) and acceptor (red) fluorescence emission, and FRET efficiency (*E_FRET_*, blue) for chromatin fibers containing *S2* at the indicated KCl and Rap1 concentrations. All Rap1 experiments were performed at 150 mM KCl. **d**) Similar to **c**) but for chromatin lacking Rap1 binding sites (*NS*). **e**) Histograms of *E_FRET_* of *S2*-containing chromatin fibers at the indicated KCl and Rap1 concentrations. All Rap1 experiments were performed at 150 mM KCl. Histograms were fitted by Gaussian functions, revealing a low-FRET (grey), medium-FRET (green) and high-FRET (red) population. Error bars are s.e.m., for the number of traces and parameters of Gaussian fits see **Supplementary Table 5. f**) Similar to **e**) but for chromatin lacking Rap1 binding sites (*NS*). **g**) Percentage of each FRET sub-population, low-FRET (LF), medium-FRET (MF) and high-FRET (HF) for chromatin containing *S2*. Box: 25-75 percentiles, whiskers: outliers (factor 1.5), line: median, open symbol: mean. For number of experiments see **Supplementary Table 5. h**) Similar to **g**) but for chromatin lacking Rap1 binding sites (*NS*). **i**) Percentage of dynamic traces for *S2* and *NS* chromatin. Box: similar to **h**). For the identification of dynamic traces see **Materials & Methods**).

First, we characterized the conformations exhibited by these chromatin fibers using FRET as a function of ionic strength. Importantly, we used two fiber types, either carrying a Rap1 site (*S2*) or without any target site (*NS,* as a control). Reconstituted chromatin fibers were immobilized in a flow channel and movies were recorded under smTIRF conditions (**Fig. 5b**). From the resulting time traces (**Fig. 5c,d**) we constructed FRET histograms (reporting on local chromatin conformation), which were fitted with 3 Gaussian functions (**Fig. 5e,f**). Under low salt conditions (40 mM KCl), we observed a population of chromatin fibers that exhibited medium FRET (MF) (E_FRET_ ∼ 0.3), corresponding to open or dynamic chromatin fibers, as well as a minor low FRET (LF) population (E_FRET_ < 0.1) (**Fig. 5c-f**). Upon raising the ionic strength to native levels (150 mM KCl), the MF population was depleted and a high FRET (HF) population appeared (E_FRET_ ∼ 0.5), indicating chromatin compaction and nucleosome stacking (**Fig. 5c-f**, **Supplementary Tables 4**). Of note, the addition of divalent cations (4 mM Mg^2+^) resulted in a similar increase in FRET efficiency (E_FRET_ of HF ∼ 0.6, **Supplementary Fig. 12**).

We could now probe the effect of Rap1 invasion on the conformational ensemble of these chromatin fibers. We thus titrated Rap1 (lacking any fluorescent label) to the chromatin fibers with (*S2*) or without (*NS*) a Rap1 binding site, using concentrations from 50 – 500 pM. The Rap1-binding site containing chromatin fibers showed a conformational response: with increasing Rap1 concentration, the fraction of tightly compacted chromatin (the HF population) disappeared, and locally opened chromatin (MF) was populated (**Fig. 5e,g** and **Supplementary Table 5**). In contrast, chromatin lacking Rap1 binding sites was not sensitive to Rap1 addition (**Fig. 5f,h**). A subset of traces (∼18-25%) exhibited anticorrelated fluctuations in the donor- and acceptor fluorescence channels, indicative of conformational dynamics in the seconds time-scale (**Supplementary Fig. 12j**). Rap1 dependent chromatin remodeling for *S2*, but not for *NS*, was associated by an increase in dynamic traces (31-37%), indicating an increase in chromatin dynamics (**Fig. 2i**). This directly indicates that Rap1 samples compact chromatin, and invades chromatin structure, most probably by exploiting intrinsic chromatin fiber dynamics. Once bound, local higher-order structure is disrupted by the pTF, thereby enabling chromatin access for subsequent TFs or the transcription machinery.

### Rap1 targets nucleosomes for eviction by RSC

Taken together, our biophysical analyses show that Rap1 increases accessibility within compact chromatin fibers but does not, by itself, evict the bound nucleosomes. Importantly, this is also true for nucleosomes containing the native *RPL30* sequence: when we incubated *RPL30* nucleosomes with Rap1, their integrity was not compromised (**Supplementary Fig. 13**). Thus, another factor is required to explain the Rap1-dependent nucleosome destabilization observed *in vivo*, e.g. an ATP-dependent chromatin remodeler. In yeast, the RSC complex is involved in the formation and maintenance of nucleosome-free regions within promoters^27-33^, and plays an important role in the organization of ribosomal protein gene promoters as well as those of many other growth-related genes^32^. We therefore hypothesized that Rap1 modulates RSC remodeling activity at promoter nucleosomes, leading to nucleosome destabilization.

To test this hypothesis, we assembled nucleosomes containing both Rap1 binding sites *S1* and *S2*. We then used purified RSC complex and yNap1 (a nucleosome chaperone)^51^ to perform remodeling assays^52, 53^ in the presence or absence of Rap1 (**Fig. 6a**). In the absence of Rap1, RSC exhibited efficient nucleosome sliding activity as judged from native gel analysis (**Fig. 6b** and **Supplementary Fig. 14**). However, we did not detect any nucleosome eviction as the additive intensity of all nucleosome bands remained unchanged over the remodeling assay. In contrast, in the presence of Rap1 we were surprised to find that nucleosome eviction became prominent. A majority of slid nucleosomes were evicted after 90 min as judged from a strong reduction in the intensity of all nucleosomal bands on a native gel (**Fig. 6c**). Instead, we observed the appearance of a new species, corresponding to Rap1 binding to free DNA (**Fig. 6c** and **Supplementary Fig. 14d**). These results therefore indicate that Rap1-bound nucleosomes alter RSC activity, biasing its function towards nucleosome eviction.

**Figure 6:**
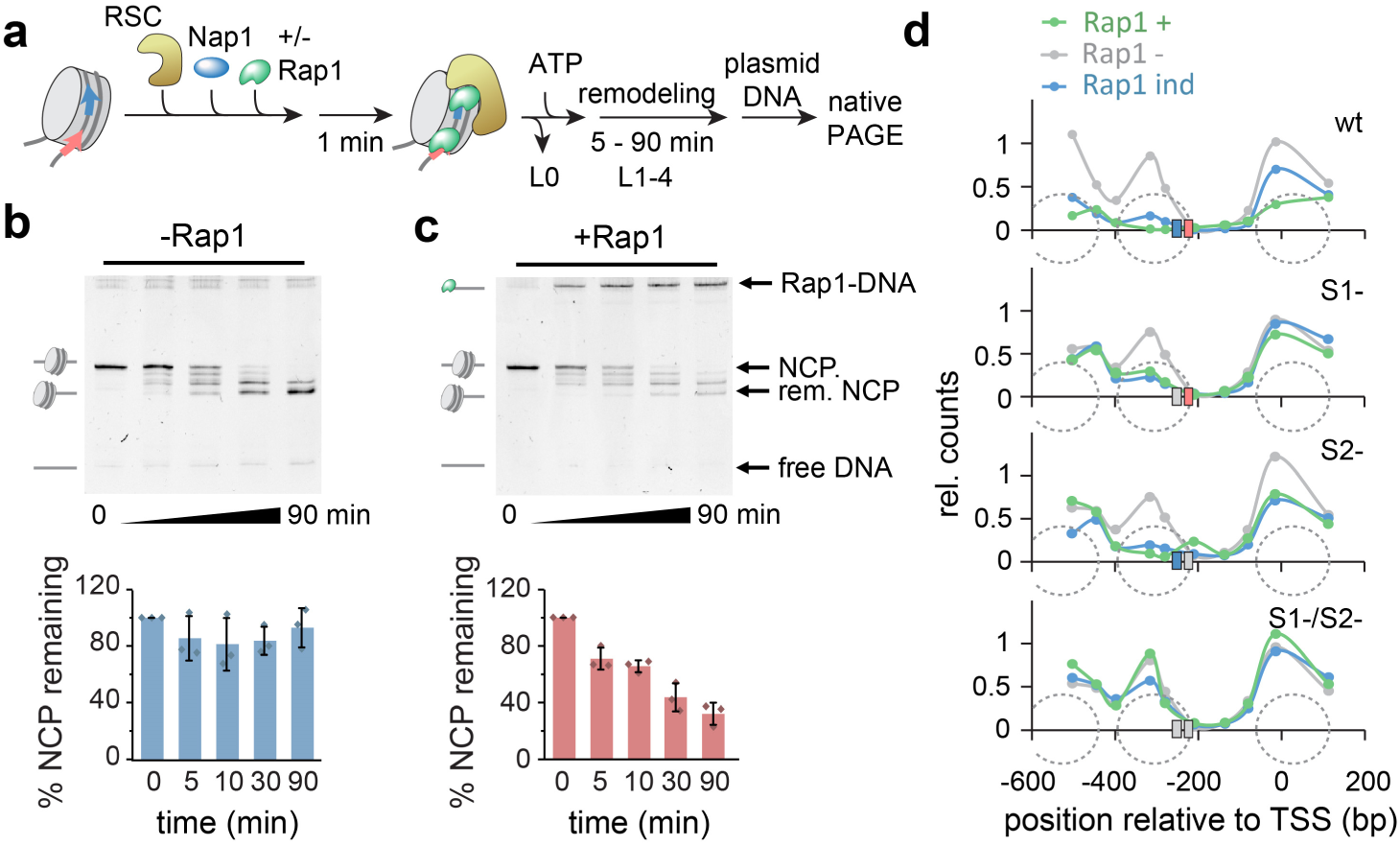
Nucleosomes marked by Rap1 are evicted by RSC. **a**) Scheme of RSC remodeling of Rap1 bound nucleosomes (containing both Rap1 binding sites1 and site2): RSC, nucleosomes and Nap1 (10eq), +/- Rap1 are combined and incubated for 1min. Then ATP is added and samples are removed at indicated time points. Each sample is incubated with plasmid DNA (which does not enter the gel) to remove RSC and Rap1, and analyzed by gel-electrophoresis as indicated (lanes L0-L4). **b**) Remodeling assays in the absence of Rap1. The bar graph below indicates the total intensity of the nucleosomal bands. (n = 3, error bars s.d., for further experimental repeats see **Supplementary Fig. 14**). **c**) Remodeling assays with 10 eq. of Rap1. Bar graphs are integrated nucleosome bands (n = 3, error bars s.d.) **d**) Effect of Rap1 binding on nucleosome stability in the *RPL30* promoter in yeast. Nucleosome positions are determined using qPCR after MNase digestion of chromatin. Analyzed are promoters containing both Rap1 binding sites (wt), with *S1* mutated (S1-), *S2* mutated (S2-) or both binding sites mutated (S1-/S2-). Shown are data for cells, where Rap1 is present (Rap1+, green), Rap1 has been depleted from the nucleus for 1 h by anchor-away (Rap1-), and has been re-introduced for 2h by expressing a Rap1 construct from an inducible promoters (Rap1 ind, blue).

We further analyzed if such dynamic nucleosome destabilization can be observed in living yeast. We generated a yeast strain carrying a chromatinized reporter plasmid bearing the *RPL30* promoter. Nucleosome presence and positioning on this test promoter was probed by chromatin digestion using MNase, followed by quantification of the fragments using qPCR^10^. If at least one Rap1 binding site was present, Rap1 was stably bound, the −1 promoter nucleosome was destabilized (**Figure 6d**) and the reporter gene was expressed (**Supplementary Fig. 15**). In contrast, if both Rap1 binding sites carried mutations that disrupt Rap1 binding, a nucleosome residing in the NFR was detected and reporter gene expression was abolished (**Fig. 6d** and **Supplementary Fig. 15**). Interestingly, when Rap1 was depleted by an “anchor-away” approach^25, 54^, the −1 nucleosome was restored for all promoters within 1 h (**Fig. 6d**). Subsequent re-induction of Rap1 finally resulted again in rapid nucleosome loss (**Fig. 6d**). Together, this demonstrates that Rap1 plays a central role in dynamically altering local chromatin environment and the stability of bound nucleosomes. In the presence of remodeling factors, these nucleosomes are then identified and destabilized. Together, this then allows Rap1 to bind DNA tightly, thereby controlling the expression of nearby genes.

## CONCLUSIONS

Elucidating the mechanism of pTF chromatin invasion and remodeling is important for a detailed understanding of gene regulation dynamics. Here, we directly observed the chromatin invasion process of an essential yeast pTF, Rap1, using highly defined reconstituted chromatin systems. The experiments enabled us to draw the following main conclusions. First, Rap1 can bind to both nucleosomes and compact chromatin fibers, but its local dwell times are greatly reduced by higher-level chromatin organization. In contrast, for the binding sites that we probed, target search kinetics driven through nonspecific DNA interactions were not affected by chromatin structure.

Second, we found that Rap1 can access its binding sites *S1* and *S2* without altering the structure of the target nucleosomes, but, interestingly, it results in local opening of chromatin fiber structure. Stacking interactions between neighboring nucleosomes are disrupted by Rap1, which in turn increases local biochemical access.

Third, in controlled *in vitro* experiments we uncovered that Rap1-bound nucleosomes are preferentially evicted by RSC, in contrast to nucleosomes which are merely shifted by RSC in the absence of Rap1. This conclusion is supported by observations in live yeast cells, where nucleosomes directly targeted by Rap1 are dynamically destabilized. Together, these data provide a comprehensive view into how the yeast pTF Rap1 locally remodels the chromatin landscape, sculpting a nucleosome-depleted region. In the following, we shall discuss several aspects of our results in the context of the current understanding of pTF function within chromatin.

### Multimodal DNA interactions guide Rap1 nucleosome invasion

Several features enable Rap1 to rapidly sample the chromatin landscape and bind to nucleosomal DNA. First the Rap1 DBD is embedded in flanking basic regions which enable nonspecific DNA binding^55^. In our single-molecule studies, we directly observed frequent short-lived interactions that are independent on the presence of a Rap1 target motif, directly reporting on the search process driven by nonspecific DNA contacts within chromatin. Second, binding to consensus sequences is very tight, ranging in affinity from low pM (e.g. in telomeric motifs^56^) to low nM for various binding motifs^57^. We indeed observe residence times in the minutes to hours time-scale in the absence of nucleosomes. Third, the Rap1 dual Myb-domains bind DNA by the insertion of a recognition helix into the major groove, a recognition mode that does not involve complete envelopment of the target DNA^23^. Indeed, we detect binding to target sites within chromatin. However, Rap1 residence times on nucleosomes are drastically reduced compared to DNA and are highly dependent on both the nature of the target site and the position of the sites relative to the nucleosome. The reduction in dwell times, and thus in affinity, arises from a combination of partial binding site occlusion (due to DNA masking by histone contacts), as well as from the highly bent DNA structure on the nucleosome. These are both known mechanisms that affect TF affinity and sequence specificity^58^. Importantly, in comparison to other TFs, which have been found to exhibit around 1000-fold reduction in residence times on chromatin substrates^3^, Rap1 still shows significant binding at the pM concentrations probed in our assays. This is consistent with the role of Rap1 as a pioneer and based on the fact that the Rap1 DBD can engage DNA on the nucleosome structure.

In the *RPL30* promoter, the high affinity site *S1* is positioned significantly further within the nucleosome sequence compared to *S2*. Intriguingly, in a chromatin context, the observed dwell times of Rap1 are similar at both sites, due to a compensation of intrinsic target site affinity and chromatin accessibility. It remains to be seen if such compensatory effects are prevalent at other promoter types.

### Rap1 passively alters local chromatin structure

Higher-order chromatin structure reduces Rap1 dwell times but does not preclude binding. Chromatin fibers are conformationally heterogeneous, as exemplified by structural studies^59-61^ or crosslinking experiments^62^. We and others have previously shown that chromatin fiber contacts are highly dynamic^17, 20, 63^. Importantly, the basic units of chromatin organization, tetranucleosome units, transiently open, close and rearrange on a millisecond timescale^20^. This exposes all internal DNA sites over time, yielding opportunities for protein factors to gain access to even compact chromatin structure. Here, our data suggest that such local dynamic modes enable Rap1, which is retained on chromatin through nonspecific DNA interactions, to access its target sites. When bound, we find that Rap1 does not induce structural rearrangements in the nucleosome for the positions probed.

Earlier experiments based on endonuclease digestion of chromatin fibers indicated that pTFs can increase local chromatin access^5^. Here, we directly observe chromatin fiber structure as a function of pTF invasion using smFRET between neighboring nucleosomes. This provides a direct molecular view on pTF function: indeed, we find that Rap1 binding results in a local opening of chromatin fiber structure. Mechanistically, our results suggest that Rap1 can bind target sites within chromatin fibers exploiting intrinsic structural fiber fluctuations. When bound, Rap1 however reduces or completely blocks the reformation of a closed tetranucleosome unit. This not only increases the accessibility for other TFs but also enables binding of remodeling factors that establish a nucleosome-depleted region at promoters.

### Rap1 marks promoter nucleosomes for eviction

Extended nucleosome depleted regions are a key feature of active yeast promoters. Rap1 is a key driver of nucleosome destabilization or eviction^10, 64, 65^. Chromatin opening in vivo has been shown to require the Rap1 DNA binding domain^64^, and it is independent on subsequent TFs also found in Rap1 regulated promoters^65^. Here, we report that Rap1, by itself, is not sufficient to disrupt targeted nucleosomes, but that it can act together with a remodeling factor, RSC, to achieve nucleosome eviction. RSC is efficient in remodeling nucleosomes, independent of Rap1^53^. However, in the presence of Rap1, nucleosome loss is accompanied by the accumulation of naked DNA containing bound Rap1 molecules. An attractive mechanistic model for this observation is that Rap1 might act as a “backstop” for RSC, inhibiting backsliding of a nucleosome over Rap1-bound DNA. Alternatively, the very high binding energy of Rap1 to free DNA (as opposed to the looser association of Rap1 with nucleosomes) might result in the local distortion of the nucleosome structure when RSC slides the histone octamer back onto Rap1-bound DNA, resulting in partial histone loss. This model (**Figure 7**) is consistent with recent observations of the collaboration between RSC and general TFs in yeast^32, 33^ and provides an explanation for the observation of MNase sensitive fragile nucleosomes at Rap1 bound promoters^25^. However, DNA binding site stability might play a key role in this mechanism since more weakly bound TFs are efficiently cleared by a passing nucleosome-remodeler complex^66^. Finally, upon clearing of the promoter region, Rap1 bound to free DNA sites is no longer impaired by chromatin structure, which results in the long residence times observed for specifically bound Rap1 *in vivo*^67^. Together, our studies thus provide a mechanistic view into how Rap1 accesses chromatin and establishes an active promoter conformation.

**Figure 7:**
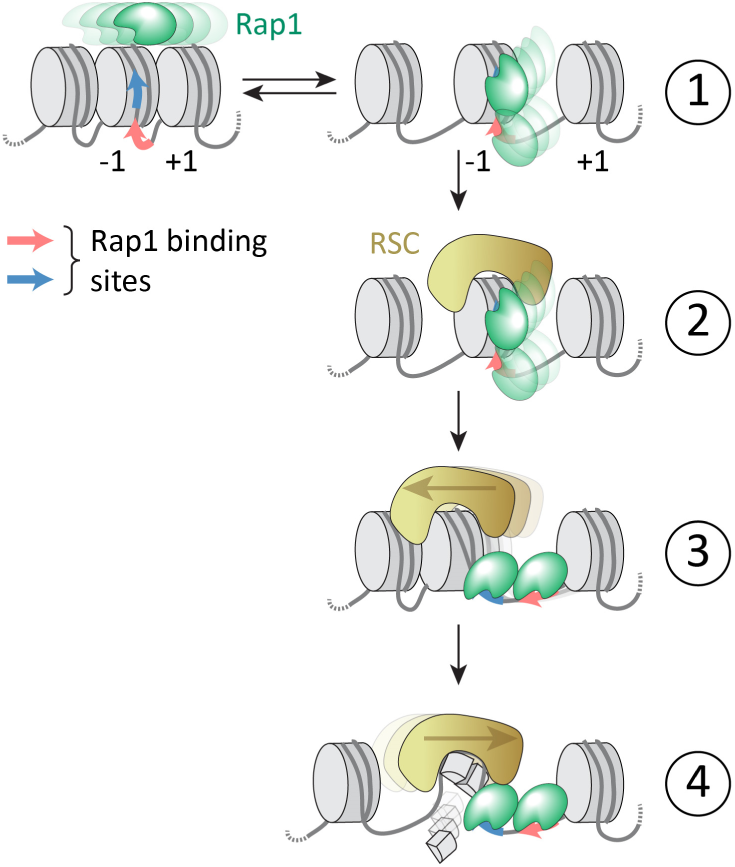
A dynamic model for Rap1 mediated promoter chromatin remodeling. Rap1 searches chromatin and its dynamic binding to a promoter site results in local chromatin opening (step 1). RSC mediated nucleosome sliding further exposes Rap1 binding sites (step 2). The fully exposed binding sites allow stable Rap1 binding (step 3). The combination of bound Rap1 and RSC remodeling results in destabilization of the −1 promoter nucleosome (step 4).

## Acknowledgements

We thank the Swiss National Science foundation (SNSF, grant 31003A_173169 to B.F. and grant 31003A 170153 to D.S.), the NCCR Chemical Biology, EPFL and the Republic and Canton of Geneva for funding. We thank Luke Laevis (Janelia Research Campus) for Halo-JF-549. We thank Louise C. Bryan for human histone octamers.

## MATERIALS & METHODS

### Expression and purification of Rap1-Halo

The Strep-MBP-TEV-Rap1-Halo construct (**Supplementary Fig. 3a**) was cloned into pACEBac1 (Geneva Biotech) and baculovirus particles were generated using the Geneva Biotech system per manufacturer’s instructions.

For Rap1 expression, 1L cultures of Sf9 cells were grown to 2 - 2.5×10^6^ cells/mL. Subsequently, the cells were infected with baculovirus, and the cultures were incubated for 3 days at 27° C, before harvesting through centrifugation (1500 rcf, 4°C for 20 min). Supernatants were discarded and pellets were resuspended in PBS, containing protease inhibitors (Roche) (10 ml PBS/L of culture), flash frozen and kept at −80°C.

For a typical purification of Rap1-Halo, 12-15 g of frozen pellets were thawed at room temperature with 36 mL of lysis buffer (200 mM KCl, 2 mM DTT, 100 mM Tris-HCl (pH 7.5), 50 mM MgOAc, 0.1% NP-40, Protease inhibitor cocktail (Roche), 1mM PMSF and 20 μl DNaseI (NEB)). Pellets, were stirred with a magnetic stir bar until fully thawed and then kept on ice. The lysate was spun for 35 min at 35000 rpm at 4°C (Ti70 rotor, Beckman Coulter) and the supernatant was filtered through a 5 µm syringe filter (Millex, Millipore). The cleared lysate was loaded onto a Strep-Trap column (GE, AKTA system), pre-equilibrated with lysis buffer. The column was washed with storage buffer (200 mM KCl, 10 mM HEPES pH 7.6, 50 mM MgOAc, and 5 mM β-mercaptoethanol (βME)) and the protein was eluted with 5 × column volumes (CV) of elution buffer (storage buffer containing 2.5 mM desthiobiotin). Fractions containing Rap1 were identified by SDS-Page (**Supplementary Fig. 3b,c**), pooled and concentrated to ∼ 500 μl total volume using Amicon 10k molecular weight cut-off (MWCO) centrifugal filters. The protein concentration was determined using UV spectroscopy. The MBP tag was subsequently removed by TEV protease digestion at 4°C (**Supplementary Fig. 3d**). For labelling, Janelia Fluor-549 HaloTag (Janelia, JF-549) was added at a protein to dye ratio of 1:1.5 followed by incubation for 1h. Labeled Rap1 was finally purified by size exclusion chromatography (SEC) using a Superose6 10/300 GL column (GE healthcare) in storage buffer using a flow-rate of 0.4 mL/min (**Supplementary Fig. 3e**). Fractions were analysed using SDS-PAGE (**Supplementary Fig. 3f**), clean fractions were pooled, concentrated (Amicon 10k MWCO filter) and protein concentrations were determined using UV spectrophotometry (at A280 and A571). Finally, labelling efficiency was calculated by using the extinction coefficients for Rap1 (107’065 mol^-1^ cm^-1^) and JF-549 (101’000 mol^-1^ cm^-1^). Typical labelling efficiency was found to be >90%.

### Expression and purification of recombinant histones

Histones were expressed and purified as described in ref. ^1^. Briefly, individual *wild-type* human histones were cloned into pet15b plasmid vectors and expressed in BL21 DE3 plysS cells. Cells were grown in LB media containing 100 µg/mL ampicillin and 35 µg/mL chloramphenicol at 37°C until the OD600 reached 0.6. Expression was induced by IPTG addition to a final concentration of 0.5 mM. After 3 h expression, cells were harvested by centrifugation and resuspended in lysis buffer (20 mM Tris pH 7.5, 1 mM EDTA, 200 mM NaCl, 1 mM βMe, Roche protease inhibitor) and frozen. Cells were lysed by freeze-thawing and sonication. Inclusion bodies were harvested by centrifugation. The inclusion body pellet was washed once with 7.5 mL of lysis buffer containing 1% triton and once without. Inclusion body pellets were resolubilized in resolubilization buffer (6 M GdmCl, 20 mM Tris pH 7.5, 1 mM EDTA, 1 mM βMe) and dialyzed into urea buffer (7 M urea, 10 mM Tris, 1 mM EDTA, 0.1 M NaCl, 5 mM 1 mM βMe, pH 7.5). Histones were purified by cation exchange chromatography using a HiTrap SP HP 5 mL column. Fractions were analyzed by SDS-PAGE and pooled, followed by dialysis into water and lyophilization. Final purification was performed by preparative RP-HPLC. Purified histones were lyophilized and stored at −20°C until used for octamer refolding.

### Large scale generation of recombinant plasmids

Plasmids containing recombinant DNA fragments for chromatin DNA assembly, which have been prepared previously^2^ (**recP1**, **recP5**) or were newly generated using restriction digestion and ligation of previous fragments (**recP1P2** or **recP4P5**, **Supplementary Fig. 6b**) were transformed into DH5α cells (for sequence information see **Supplementary Table 1**). Cells were cultured overnight in 6L 2xTY medium and harvested by centrifugation. For alkaline lysis, the cells were resuspended in 120 mL lysis solution I (50 mM glucose, 25 mM Tris pH 8, 10 mM EDTA). 240 mL lysis solution II (0.3 M NaOH, 1% SDS) was added and mixed by stirring. 240 mL lysis solution III (4 M KAc, 2 M acetic acid) was added to neutralize the solution which was left at 4° C for 15 min. After centrifugation, the supernatant was passed through Miracloth (Merck). Plasmid DNA was collected by isopropanol precipitation: 0.52 volume equivalents of isopropanol was added followed by centrifugation at 11’000 × g for 20 min at 4° C. The DNA pellet was dissolved in TE 10/50 (10 mM Tris pH 7.5, 50 mM EDTA) in the presence of 100 units of RNAse A, and digested for 2 h at 37° C. To perform SEC the buffer was adjusted to 2M KCl (10 mM Tris, 50 mM EDTA and 2 M KCl). The plasmid was then purified in the same buffer on a XK 50/30 column (GE Healthcare) containing a bed of 550 mL sepharose 6 Fast Flow (GE Healthcare). Eluted plasmid DNA from was precipitated with isopropanol. The pellet was finally dissolved in TE 10 / 0.1 (10 mM Tris pH 7.5, 0.1 mM EDTA) and stored at −20° C.

### Large scale restriction digest and purification of recombinant plasmids

Purified plasmid DNA was collected by isopropanol precipitation and the DNA pellet was dissolved in milliQ H_2_O. For a typical reaction, either 200 units of DraIII-HF (NEB) (for **recP1P2**) or 200 units of BsaI-HF (NEB) (for **recP4P5**) or 200 units of both DraIII-HF and BsaI-HF (NEB) (for **recP1** and **recP5**) were added to 200 pmol of plasmid DNA in 200µl 1x NEB CutSmart buffer. After 8-10 h digestion at 37°C, digestion progress was analyzed by gel electrophoresis on a 1% agarose gel (run in 1 × TBE running buffer, 100 V, for 50 mins) to check completeness (**Supplementary Fig. 6c**). If required, the digestion was pushed to completion by adding another 100 units of enzyme and incubation for further 8-10 h at 37° C. Once the digestion was complete, 100 units EcoRV-HF (NEB) was added and left 8-10 h at 37°C. Complete digestion was verified by electrophoresis as described above (**Supplementary Fig. 6c**). If the digestion was not complete an additional 50 units of enzyme was added and left 8-10 h at 37° C. Once the digestion was complete, the desired chromatin DNA fragments were purified from the plasmid remnants through successive PEG precipitations. This involves adding 40% PEG 6000 to the digestion reactions until a final concentration of 5-6% PEG 6000 was reached. Additionally, the NaCl concentration was adjusted to 0.5 M. The sample was then spun at 20’000 × g at 4° C for 20 min. The supernatant was collected, and PEG 6000 was added to the supernatant to increase the final PEG % by increments of 0.5 %. The sample was then spun at 20’000 × g at 4° C for 20 min. This was repeated until a suitable purity was achieved (**Supplementary Fig. 6d**). Finally, the chromatin DNA fragments were isolated using QIAquick PCR purification spin columns (Qiagen).

### Oligonucleotide labelling

Fluorescently labeled oligonucleotides were generated as described in ref. ^2^. Briefly, 5-10 nmol of single stranded oligonucleotide, containing amino modified C6 dT, was diluted in 25 µl 0.1 M sodium tetraborate, pH 8.5. 5 µl of a 5 mM stock of succinimidyl-ester modified fluorophore (Alexa 568, Alexa 647 or Cy3B) were added to the reaction mix and left shaking at room temperature for 4 – 8 hours. For a table enumerating all labeled oligonucleotides see **Supplementary Table 2**.

Reaction progress was analyzed by RP-HPLC using a gradient from solvent A (95% 0.1M triethylammonium acetate (TEAA) pH 7, 5% ACN) to solvent B (70% 0.1M TEAA pH 7, 30% ACN) on a 3 µm 4.6×150 mm InertSustain C18 column (GL sciences) over 20 min. More dye was added when required. For purification, the labeled DNA was ethanol precipitated (by the addition of 2.75 equivalents of cold ethanol, 0.3M NaOAc pH 5.2, followed by centrifugation at 20’000 × g at 4° C for 20 min) twice successively to remove excess unconjugated dye. The DNA pellet was finally dissolved in 100 µl solvent A and purified by HPLC. The purified DNA was finally ethanol precipitated and dissolved in milliQ water to a concentration of 2.5 µM.

### Production of labelled DNA fragments

Labeled DNA was prepared by PCR (fragments **P2, P3_S1**, **P3_S2**, **P3_S2\***, **P3_S1S2** and **P4**, for sequences and labeling schemes, see **Supplementary Table 1-3**). For a typical reaction, 96 × 50 ul PCR reactions in 1 × ThermoPol reaction buffer (NEB) were prepared using template (0.01 ng µL^-1^), forward primer (0.250 µM), reverse primer (0.250 µM), dNTPs (0.2 mM, NEB) and Taq DNA polymerase (1.25 units, NEB). A typical program included an initial step of 12 s at 94° C, followed by 30 cycles of 12 s at 94° C, 12 s annealing at 58-65° C and 12 s extension at 72° C. Final extension was also done at 72° C for 12 s. PCR reactions were subsequently purified using QIAquick PCR purification spin columns (Qiagen) (**Supplementary Fig. 2a**).

About 0.33 nmol of PCR generated DNA (**P3_S1**, **P3_S2**, **P3_S2\***, **P3_S1S2** and **P3_Rpl30, Supplementary Table 1**) was digested in 200 µl of 1 × CutSmart buffer using 100 units of BsaI-HF (NEB) and 100 units of DraIII-HF (NEB) for 8-10h at 37° C. The progress of the digestion was analyzed on a 2% agarose gel (running conditions: 1 × TBE, 110 V for 50 min) (**Supplementary Fig. 2b**). Finally the DNA fragments were purified using QIAquick PCR purification spin columns (Qiagen) and the concentration was determined by UV spectroscopy.

### Ligation and purification of 1 × 601 DNA to biotin anchor

For the generation of nucleosome DNA for single-molecule experiments, a biotin containing anchor (**Anchor_rev, Supplementary Table 2**) was annealed to its complementary strand containing a phosphorylated 5’- BsaI overhang (**P3_Anchor_fwd**, **Supplementary Table 2)** and a 10-fold excess was added to 150-300 pmol (∼20-40 µg) of digested PCR generated DNA (**P3_S1**, **P3_S2**, **P3_S2\*** and **P3_S1S2)** in 100 µl 1x T4 ligase buffer (NEB). Upon complete ligation of digested DNA, excess biotin anchor was removed by PEG precipitation (**Supplementary Fig. 2c-d**). Finally, the DNA fragments were purified using QIAquick PCR purification spin columns (Qiagen) and the concentration was determined by UV spectroscopy.

### Mononucleosome nucleosome formation

Nucleosomes (MN_S1, MN_S2, MN_S2*, MN_S1S2, MN_S1_FRET, MN_S2_FRET, MN_Rpl30, **Supplementary Table 2**) were prepared following ref. ^3^. Typically, 1-5 µg of labelled and biotinylated DNA (**P3_S1**, **P3_S2**, **P3_S2\*** and **P3_S1S2**) was combined with purified refolded octamers at experimentally determined ratios (1:1 to 1:2, DNA to histone octamer) in 10 µl TE (10 mM Tris-HCl pH 7.5, 1 mM EDTA) supplemented with 2 M KCl. After a 30 min incubation period at room temperature, 10 µl TE was added and further incubated for 1 h. This was followed by sequential addition of 5 µl TE, 5 µl TE and finally 70 µl TE with 1 h incubation periods in between each addition, to arrive to 0.2 M KCl. Samples were then spun at 20’000 × g for 10 min at 4 °C and the supernatant was kept on ice. To determine the quality of mononucleosome assemblies, 0.8% Agarose 0.25 × TB gels were run at 90 V on ice for 90 min (**Supplementary Fig. 2f-I** and **Supplementary Fig. 13b-c**).

### Electrophoretic mobility shift assays (EMSA)

EMSAs to determine Rap1 binding to DNA were done in single-molecule imaging buffer (IB, 50 mM HEPES pH 7.5, 130 mM KCl, 10% v/v glycerol, 0.005% v/v Tween 20, 2 mM Trolox, 3.2% w/v glucose), in the presence of 50 ng/ul poly-d(I-C) (Roche) and with 20 ul total volume. Typically, 200 nmol stocks of DNA and 3 µM stocks of Rap1-Halo were prepared and serially diluted to desired concentrations. Reactions were mixed by pipetting and left for 10 min at room temperature. Sucrose was added to a final concentration of 8% and reactions were loaded onto 5% Polyacrylamide gels run in 0.5 × TBE at 100 V for 60 min. Images were taken using ChemiDoc MP (Biorad) (**Supplementary Fig. 1d,e**). For densitometry quantifications, ImageLab (Biorad) software was used for band quantification of bound and unbound fraction of DNA. The data was analyzed in Origin (OriginLab) by non-linear curve-fitting using a sigmoidal function to determine Kd (**Supplementary Fig. 1 d,e**).

### Convergent 3-piece and 5-piece convergent DNA ligation for synthesis of 12×601 DNA

Singly-labeled and biotinylated 12×601 DNA was produced as shown in **Supplementary Figure 6a**. Typically, 50-60 pmol of PEG purified restriction enzyme digested **recP1P2** and an excess 1×601 P3 (**P3_S1** and **P3_S2)** (between 20-30 % excess) were added to 200 μl 1 × T4 ligase buffer containing 400 units of T4 ligase. The reaction was followed using 1% agarose gels (**Supplementary Fig. 6e**). Upon completion, **P1P2P3** was PEG purified (**Supplementary Fig. 6f**) and added to excess **P4P5** and biotin labelled anchor (20-30% excess **P4P5** and 10-fold excess biotin anchor). The reaction was followed using 1% Agarose gels (**Supplementary Fig. 6g)**. Upon completion, the complete DNA **P1P2P3P4P5A** was PEG purified and subsequently purified using Qiaquick PCR purification spin columns (Qiagen), the concentration was determined by UV spectrophotometer (**Supplementary Fig. 6h-i**).

The FRET pair Cy3B and Alexa647, were site-specifically introduced respectively on **P2** and **P4** at the 39-base-pair position relative to the dyad in the 601 sequence (**Supplementary Figure 11c-d**). About 30 pmol of each piece was used for 5-piece convergent DNA ligation to produce two intermediate 6 × 601 pieces as followed: **recP1** was ligated to Cy3B-labeled **P2** in 20% excess for 2 h using T4 DNA ligase, then unlabeled **P3** in 20% excess relative to **P2** was added and left to ligate another 15 h. Similarly, **recP5** was ligated to 20% excess Alexa647-labeled **P4** for 15 h (**Supplementary Figure 10f**). Singly-labeled 6×601 intermediate fragments **P1-3** and **P4-5** were PEG purified from individual pieces. Pellets containing enriched fragments were collected, dissolved in 50 µL TE buffer (10 mM Tris, 0.1 mM EDTA, pH 8.0), and used for the final ligation (**Supplementary Figure 11g**). A biotinylated anchor was added into the final ligation of 2 intermediate 6×601, and the reaction was proceeded for 15 h at room temperature. PEG precipitation was performed similarly to previous step, and the enriched final products were collected and purified using Qiaquick PCR purification spin columns (Qiagen), the concentration was determined by UV spectrophotometer (**Supplementary Figure 11a-c**).

### Reconstitution of 12-mer chromatin fibers

Chromatin fibers (CH_S1, CH_S2, CH_S1_FRET and CH_S2_FRET, **Supplementary Table 3**) were reconstituted from singly/doubly-labeled and biotinylated 12×601 DNA and wild-type recombinantly purified human histone octamers. In a typical dialysis, 200-300 pM 12×601 DNA, 0.5-1 equivalents of MMTV DNA and reconstituted octamers (using experimentally determined DNA:octamer ratios) were added to a micro-dialysis unit (Thermo Scientific, Slide-A-Lyzer – 10’000 MWCO), then dialyzed in TE buffer (10 mM Tris pH 7.5, 0.1 mM EDTA pH 8.0) with a linear gradient from 2 M to 10 mM KCl for 16-18 h, and finally kept in TEK10 buffer (10 mM Tris pH 7.5, 0.1 mM EDTA pH 8.0, 10 mM KCl) for another 1 h. Chromatin assemblies were centrifuged at 21’000 × g for 10 min at 4° C, the supernatant was then transferred to a fresh tube. The concentration and volume of the chromatin assemblies was determined using UV spectrophotometer. Chromatin assembly quality was controlled by the appearance of MMTV nucleosomes and ScaI digestion of 12x assemblies. Digestion reactions were analyzed on a 0.8% agarose gel and 5% TBE polyacrylamide gel electrophoresis. All experiments were carried out at 4° C (**Supplementary Figures 7** & **11**).

### Preparation of microfluidic chambers for sm-FRET/TIRF experiments

Cleaning, silanization and PEGylation of coverslips and glass slides was done described previously in ref. ^1^. Briefly, coverslips (24 × 40 mm, 1.5 mm thickness) and glass slides (76 × 26 mm with 2 rows of 4 holes drilled) were sonicated for 20 mins in 10% Aconox, rinsed with milliQ water and the procedure repeated sequentially with acetone and ethanol. Both coverslips and glass slides were then placed in piranha etching solution (25% v/v 30% H_2_O_2_ and 75% v/v H_2_SO_4_) for minimum 2 h. After thorough washing with milliQ H_2_O, coverslips and slides were sonicated in acetone for 10 min, then incubated with 2% v/v aminopropyltriethylsilane (APTES) in acetone for 15 min, and dried. Flow-chambers were assembled from one glass slide and one coverslip separated by double-sided 0.12 mm tape (Grace Bio-labs) positioned between each hole in the glass slide, and the open ends were sealed with epoxy glue. Pipette tips were fitted in each of the 2 × 4 holes on each side of the silanized glass flow chambers as inlet reservoir and outlet sources and glued in place with epoxy glue. The glue was allowed to solidify for 30-40 min. Subsequently, 350 µL of 0.1 M tetraborate buffer at pH 8.5 was used to dissolve ∼1 mg of biotin-mPEG(5000 kDa)-SVA, and 175 µL from this was transferred to 20 mg mPEG (5000kDa)-SVA. This was centrifuged and mixed to homogeneity with a pipette before 40-45 µL aliquots were loaded into each of the four channels in the flow chamber. The PEGylation reaction was allowed to continue for the next 2½-4 h after which the solution was washed out with degassed ultra-pure water (Romil).

### Single-molecule TIRF (sm-TIRF) co-localization microscopy measurements

Measurements were done according to ref. ^1^. Objective-type smTIRF was performed using a fully automated Nikon Ti-E inverted fluorescence microscope, equipped with an ANDOR iXon EMCCD camera and a TIRF illuminator arm, controlled by NIS-elements and equipped with a CFI Apo TIRF 100x oil immersion objective (NA 1.49), resulting in a pixel size corresponding to 160 nm. Laser excitation was realized using a Coherent OBIS 640LX laser (640 nm, 40mW) and coherent OBIS 532LS laser (532 nm, 50 mW) on a custom setup laser bench. Wavelength selection and power modulation was done using an acousto-optical tunable filter (AOTF) controlled by NIS-elements. Typical laser intensities in the objective used for measurements were 0.8 mW for both 532 nm and 640 nm laser lines. For all smTIRF experiments, flow channels were washed with 500 µL degassed ultrapure water (Romil), followed by 500 µL 1 × T50 (10 mM Tris pH 8, 50 mM NaCl) and background fluorescence was recorded with both 532 nm and 640 nm excitation. 50 µL of 0.2 mg/ml neutravidin was then injected and incubated for 5 mins, and washed using 500 µL 1xT50. 50 pM of Alexa647 labelled DNA/mononuceosomes/12-mer chromatin assemblies were then flowed in for immobilization in T50 with 2 mg/ml bovine serum albumin (BSA, Carlroth) (25 × 50 µm imaging area was monitored using 640 nm excitation to check for sufficient coverage). 500 µL 1 × T50 was used to wash out unbound Alexa647 labelled DNA/mononuceosomes/12-mer chromatin assemblies. 50-100 pM JF-549 labelled Rap1-Halo (see table below for details) was flowed in using imaging buffer (50 mM HEPES pH 7.5, 130 mM KCl, 10% v/v glycerol, 0.005% v/v Tween 20, 2 mM Trolox, 3.2% w/v glucose, 1x glucose oxidase/catalase oxygen scavenging system and 2 mg/ml BSA). Images were recorded using the following parameters:

| | Camera $t_{on}$<br>(msec) | Camera $t_{off}$<br>(msec) | Orange<br>channel #<br>frames | Far-red<br>channel: #<br>frames | n repeat |
| --- | --- | --- | --- | --- | --- |
| DNA | 100 | 1400 | 1 | 1 | 5000 |
| Mono-nucleosome | 100 | 0.3 | 199 | 1 | 40 |
| 12-mer chromatin fiber | 100 | 0.3 | 199 | 1 | 40 |

Here $t_{on}$ denotes the camera integration time, whereas $t_{off}$ indicates interspersed time intervals of camera inactivity.

Each experiment was repeated several times (see **Table 1** for number of repeats), using at least two independently produced chromatin preparations on two different days.

### Photobleaching test for JF-549 Rap1-Halo

Slides were prepared as described in the preceding sections. However, no BSA was added to imaging buffer (50 mM HEPES pH 7.5, 130 mM KCl, 10% v/v glycerol, 0.005% v/v Tween 20, 2 mM Trolox, 3.2% w/v glucose, 1x glucose oxidase/catalase oxygen scavenging system). JF-549 labelled Rap1-Halo was flown into the channel and non-specifically adsorbed on the glass surface. Movies were recorded using continuous 532 nm illumination (t_on_ 50 msec and t_off_ 0.3 msec) using the indicated excitation laser powers (**Supplementary Fig. 4b**). Absolute laser power was determined using a laser power meter at the objective.

### Image processing, single-molecule trace extraction and trace analysis

Single-molecule trace extraction and trace analysis were done according to ref. ^1^ with some adjustments. Firstly, a background subtraction was performed for all Rap1-Halo binding movies using a rolling ball background subtraction in ImageJ (using 50 pixel rolling ball size). Using a custom built Matlab (Mathworks) program suite, DNA/nucleosome or chromatin positions were detected via a local maxima approach. Sequential images were aligned using the far-red channel to compensate for stage drift. Fluorescence intensities (in the orange channel) were extracted from the stack within a 2 pixel radius of the identified DNA peaks. Every detected spot in the orange channel was fitted with a 2D-Gaussian function to determine co-localization with immobilized DNA/chromatin. Peaks exceeding an experimentally determined PSF width for a single JF-549 molecule were excluded from further analysis. Extracted fluorescence traces were filtered using a forward-backward non-linear filter^4^ to reduce noise.

Residence times were determined using a semi-automatic procedure. Individual binding events were detected using a thresholding algorithm. Overlapping multiple binding events were excluded from the analysis. For each movie cumulative histograms were constructed from detect bright times (*t_bright_*) corresponding to bound Rap1 molecules, usually including data from ∼100 individual traces. The cumulative histograms from traces corresponding to individual DNA / mononucleosome / chromatin fibers were fitted with either di- or tri-exponential functions:

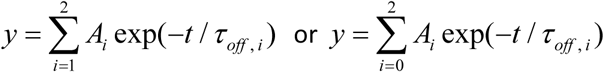

yielding non-specific residence times *r*_off,0_ or the specific residence times *r*_off,1_ and *r*_off,2_ (**Figures 2, 3** and **Supplementary Fig. 4a, c-k. Supplementary Fig. 5a-i**). Cumulative histograms constructed from dark times (*t_dark_*), in between binding events, were fitted with mono-exponential functions:

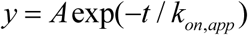

to obtain apparent on-rates. The detected on-rates contain both contributions from non-specific and specific binding events. To calculate specific on-rates (*k_on_*), the contributions from non-specific events have to be filtered out. To this end, measured *k_on_*_,app_ values were corrected using the amplitude contributions of non-specific (*A_0_*) and specific binding events (*A_1_, A_2_*).

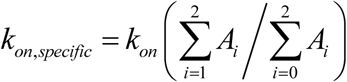

### Single-molecule FRET (smFRET) measurements for chromatin conformation analysis

#### Measurement procedure

Flow cell preparation and chromatin loading was performed as described in ref. ^2^ and the preceding paragraphs. Experiments were performed in FRET imaging buffer (40 mM KCl, 50 mM Tris, 2 mM Trolox, 2 mM nitrobenzyl alcohol (NBA), 2 mM cyclooctatetraene (COT), 10% glycerol and 3.2% glucose) supplemented with GODCAT (100x stock solution: 165 U/mL glucose oxidase, 2170 U/mL catalase). Experiments on chromatin remodeling effect of Rap1 were performed with imaging buffer containing 150 mM KCl, and 0.1 mg/ml of BSA was added to prevent non-specific binding of Rap1 to glass surface. For Rap1 titration, unlabeled Rap1-Halo was used.

smFRET data acquisition was carried out with a micro-mirror TIRF system (MadCityLabs) using Coherent Obis Laser lines at 405 nm, 488 nm, 532 nm and 640 nm, a 100x NA 1.49 Nikon CFI Apochromat TIRF objective (Nikon) as well as an iXon Ultra EMCCD camera (Andor), operated by custom-made Labview (National Instruments) software.

For general smFRET imaging, a programmed sequence was employed to switch the field of view to a new area followed by adjusting the focus. The camera (at 500 EM gain) was triggered to acquire 1950 frames with 532 nm excitation and 100 ms time-resolution followed by a final change to 640 nm excitation.

Each experiment was repeated several times (see **Supplementary Tables 4** and **5** for number of repeats), using at least two independently produced chromatin preparations on two different days.

#### Calibration

Before each experiment, instrument calibration was performed by imaging 100-nm biotinylated Gold nanoparticles (Cytodiagnostics) with 532 nm excitation and 100 ms time-resolution over 10 s. Acquired calibration movies were analyzed using a custom-written Macro ImageJ to determine the signal-to-noise ratio (*SNR*) as follows:

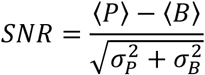

Where 〈P〉 and *a_P_* are average and standard deviation of peak, and 〈B〉 and *a_B_* are that of its background. Our standard calibration was performed with 12 mW of 532 nm excitation at 500 EM gain of the camera resulting in average *SNR* of 6.5-8.5. Moreover, at least well-separated 10 nanoparticles representing the field of view and appearing in both the donor and the acceptor channels were selected to generate a transformation matrix, which was further applied for aligning non-isotropically donor and acceptor images.

#### smFRET data analysis

FRET reporting on chromatin conformation as a function of ionic strength or Rap1 binding was recorded as described above.

For FRET calculation, the orange and far-red channel detection efficiency ratio *y* and donor dye bleed-through parameter *f3* were independently determined using double-stranded DNA oligonucleotides, where X and Y indicate respectively 5’-Amino-C6 and 5-C6-Amino-dT and labeled respectively with Cy3B and Alexa Fluor 647

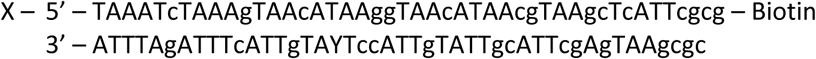

For all recorded movies, background correction was performed in ImageJ using a rolling ball algorithm. Single-molecule kinetic trace extraction and analysis was performed in custom-written MATLAB software. Donor and acceptor channels were non-isotropically aligned using the nanoparticle based transformation matrix. Individual molecules were automatically detected in the initial acceptor image prior to donor excitation, and the same peaks were selected in the donor channel. Peaks that are (*i*) tightly clustered or (*ii*) above an intensity threshold of 8000 in the donor channel and 5000 in the acceptor channels indicating aggregation or (*iii*) do not appear in both donor and acceptor channels were excluded from analysis. Kinetic donor and acceptor fluorescence traces were extracted for each single-molecule. Selection criteria were similar to ref. ^2^. Traces were included if they exhibited: (*i*) a single bleaching event, (*ii*) constant total fluorescence emission > 2000 counts from combined donor and γ-corrected acceptor channel (*iii*) a constant baseline lasting for at least 2 s after donor bleaching, (i*v*) donor emission for at least 5 s and finally (*v*) the presence of acceptor dye. The last condition is verified as follows: If the donor dye bleaches first, acceptor emission must be detectable at the end of the experiment upon direct acceptor excitation. If the acceptor dye bleaches first, a significant increase is seen in the donor channel. From selected traces, donor (*F_D_*) and acceptor (*F_A_*) fluorescence emission intensity, FRET efficiency (*E_FRET_*) was calculated as follows:

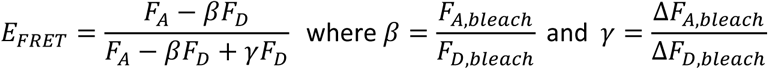

We determined the detection efficiency *y =* 0.423 and the bleed-through *f3 =* 0.073 for the FRET pair Cy3B/Alexa647 with our experimental setup. These values were used to calculate *E_FRET_* for the selected traces, and construct *E_FRET_* histograms with a bin size of 0.02. *E_FRET_* histograms of each trace of length > 5 s were normalized to total counts. Final histograms of each independent measurement were fitted using 3 Gaussian functions as follows:

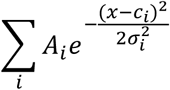

Where *A_i_* is the amplitude or the height of the fitting peak, *c_i_* is the position of the center of the peak, and *a_i_* is the standard deviation which controls the width of the Gaussian peak. The integral area of each peak was calculated as follows:

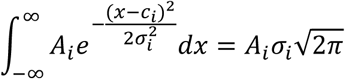

Where indicated, low-FRET (LF), medium-FRET (MF) and high-FRET (HF) refer respectively to the center of the Gaussian peak limited with *c_i_<* 0.2, 0.2 ≤ *c_i_* ≤ 0.4, and *c_i_ >* 0.4. The percentage of LF-population at compaction conditions, i.e. in high salt or presence of Mg^2+^, indicates the fraction of uncompacted chromatin, and hence reports chromatin assembly quality. Control of chromatin compaction was performed, and only measurements on chromatin preparation giving *E_FRET_* histograms with < 50% of LF-population were selected to further analysis (**Supplementary Figure 12**).

Dynamic traces were identified by fluorescence cross-correlation analysis, performed using the following function:

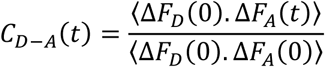

Where Δ*F_D_* and Δ*F_A_*denote the variances of donor and acceptor fluorescence at time *0* or *t*. Only traces lasting for more than 10 s, and spending > 20% of the duration time at *E_FRET_ >* 0.2 were included and fitted with a bi-exponential function. A dynamic trace is defined as the one showing a cross-correlation amplitude inferior to −0.1 and a relaxation time superior to 100 ms.

All Gaussian fit parameters and cross-correlation analysis are shown in **Supplementary Tables 4** & **5**.

### Nucleosome shift assays with RSC, Nap1 and Rap1

Purified RSC and recombinant yNap1 were used (for the purification, see ^5^). All reactions were performed in reaction buffer (10 mM Tris pH 7.4, 150 mM KCl, 3 mM MgCl_2_, 0.1 mg/ml BSA) and a total volume of 50 µl. The following components were added in sequential order mononucleosomes (to give a 20 nM final concentration), yNap1 (10 eq. yNap1 : 1 eq. mononucleosomes), if required Rap1 (10 eq. Rap1 : 1 eq. mononucleosomes), RSC complex (5 eq. RSC : 1 eq. mononucleosomes) and finally ATP (1mM). Reactions were placed at 30°C and 10µl were taken for each time point, to which was added a 3-fold excess of plasmid DNA (compared to nucleosomes) containing a Rap1 binding site and returned to 30°C for 5mins. Reactions were then placed on ice until, glucose was added to make 8% final concentration and loaded onto commercial Criterion Precast Gel (Biorad) 5% TBE, 1mm, run in 1xTBE at 200V for 35-45mins on ice. Gels were stained in Gelred and imaged using ChemiDoc MP (Biorad) (**Supplementary Fig. 14a-c**).

### Ensemble FRET measurements

All measurements were performed using a Fluorolog®-3 Horiba Jobin Yvon spectrofluorometer, in T50 buffer (10 mM Tris pH 8, 50 mM NaCl) 60 µl total volume. Nucleosomes (final concentration of 25-30 nM) and Rap1 (0, 1, 2, 5, 10 equivalents) were mixed by pipetting in T50 buffer and left for 10 min room temperature to bind. Fluorescence emission spectra are taken from 585 nm to 700 nm (1 nm increments) using 578 nm as excitation wavelength. Spectra for DNA only, T50 only and donor only samples were taken. For a given sample, NaCl was added to 800 mM to observe nucleosome disassembly. FRET efficiency was calculated from donor emission:

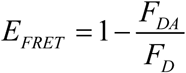

with *F_DA_* denoting donor emission in the presence of acceptor, and *F_D_* denoting donor emission in the donor-only sample. Additionally, reactions were loaded onto 0.5xTBE 5% polyacrylamide gels to check binding.

### Yeast experiments

#### Plasmid construction

The pRS313-GALL plasmid was constructed by subcloning of *Sac*I and *Xba*I fragment from pRS416-GALL plasmid and inserted into pRS313 for construction of plasmid expressing RAP1 under the control of GALL promoter. The *RAP1* coding region was amplified using primers 5’-CATGTCTAGAATGTCTAGTCCAGATGATTTTGAAAC-3’ (Forward) and 5’-CATGCCCGGGTCATAACAGGTCCTTCTCAAAAAATC-3’ (Reverse) containing *Xba*I and *Sma*I sites and inserted into pRS313-GALL construct, digested with *Xba*I and *Sma*I. To construct pLR10-RPL30 plasmids, first RPL30 WT and RPL30-m1, RPL30-m2, and RPL30-m1/m2 mutants were cloned into pUC18 plasmid between *Sph*I and *Sac*I sites using primers 5’-ATGCGCATGCCTGCGTATATTGATTAATTGAA-3’ (Forward) and 5’-ATGCGAGC TCATATCATGCAGTACATTGACAGTATATCA-3’ (Reverse). Corresponding regions were then amplified by PCR using primers 5’-ATGCGTCGACATATCATGCAGTACATTGACAGTATATCA-3’ (Forward) and 5’-ATGCGCATGCCTGCGTATATTGATTAATTGAA-3’ (Reverse), and cloned into pLR10 plasmid just upstream of the YFP reporter gene at *Sph*I and *Sal*I sites. The yeast RAP1 anchor away strain HHY168 *RAP1(1-134)-FRB1-RAP1(136-827)-LEU2* (YJB26) was co-transformed with the pRS415-GALL-RAP1 and pLR10-RPL30 plasmids.

#### Yeast growth conditions

The yeast cells, transformed with pRS313-GALL-RAP1 and pLR10-RPL30 plasmids, were grown overnight in SC-His-Ura containing 2% raffinose. Overnight cultures were diluted to OD_600_ 0.1, grown at 30°C to OD_600_ 0.3-0.4, and then treated with either vehicle (90% ethanol/10% Tween) or, for anchor-away, with rapamycin (1 mg/ml of 90% ethanol/10% Tween stock solution) at a final concentration of 1 μg/ml^6^ (1μg/mL) for 1 hr to deplete FRB-tagged RAP1 protein. Following the rapamycin treatment, the strains were grown in medium containing 2% galactose for 2 hr to induce expression of RAP1 or 2% raffinose.

#### MNase digestion and nucleosome mapping

MNase digestion was performed as described^7^. Briefly, yeast cells were grown at 30°C for o/n in SC-His-Ura media containing 2% raffinose to OD_600_ 0.3-0.4, crosslinked with 1% formaldehyde for 5 min and quenched by the addition of 125 mM glycine for 5 min at room temperature. The cell pellets were resuspended in spheroplasting buffer (1M sorbitol, 1 mM β-mercaptoethanol, 10 mg/mL zymolyase) after harvesting and incubated for 8 min at room temperature. Spheroplasts were washed twice using 1 mL of 1 M sorbitol and treated with different concentrations of MNase, ranging from 0.05 to 1.0 units. The samples were incubated at 37° C for 45 min in MNase digestion buffer (1M Sorbitol, 50 mM NaCl, 10 mM Tris pH 7.4, 5 mM MgCl_2_, 1 mM CaCl_2_, 1mM β-mercaptoethanol, 0.5 mM spermidine and 0.075% NP-40). Digestion reactions were stopped by the addition of EDTA (30 mM), the crosslinks were reversed with SDS (0.5%) and proteinase K (0.5 mg/mL) and incubated at 37° C for 1 h and then transferred to 65° C for at least 2 h. The DNA was isolated by phenol/chloroform/isoamyl alcohol (25:24:1) extraction, concentrated with ethanol and treated with RNase at 37° C for 1 h for monitoring on agarose gel (2%). MNase profiles were determined by qPCR of chromatin samples (previously digested with 0.5 units MNase) using a set of nested primer pairs covering the *RPL30* promoter region ∼561 bp upstream from the ATG.

#### Flow cytometry

Flow cytometry analysis was performed to detect the expression of a YFP reporter driven by RPL30 promoter and its variants in different conditions. Yeast transformants were grown to stationary phase overnight in appropriate media, the cells were diluted to OD600 0.1 the next day and grown to exponential phase at OD600 0.3-0.4. Upon flow cytometry, the cells were diluted 10-fold into SC-His-Ura media and immediately processed on Beckman Coulter Gallios Flow Cytometer. YFP-expressing cells were sorted by fluorescence-activated cell sorting (FACS) analyses using excitation lasers at 488 nm, and filtering emissions at 525 nm.

#### Data availability

Microscopy data, evaluation scripts and detailed plasmid maps of expression vectors are available upon request.

## SUPPLEMENTARY FIGURES & TABLES

**Supplementary Fig. 1.**
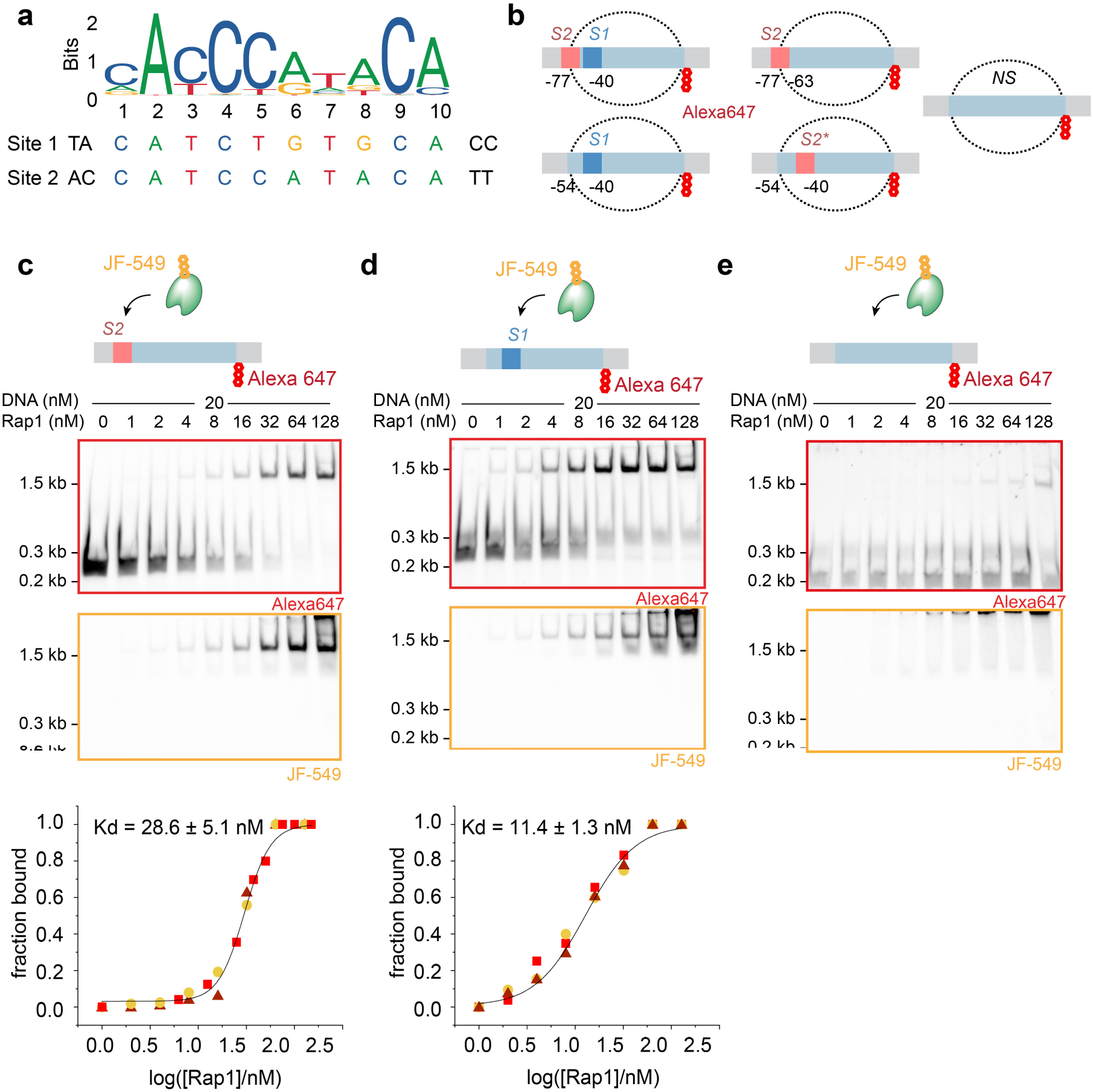
Mononucleosome design: **a)** Rap1 consensus sequence. Aligned below are the sequences of the *RPL30* promotor Site 1 (*S1*) and Site 2 (*S2*). **b)** Schematic representation of the DNA constructs used (see **Supplementary Table 1-3** for sequence information); indicated are the position of the dye and the positions of the different binding sites relative to the dyad, additionally the position of the octamer is depicted by the dotted black line. Linker regions are shown in grey. **c-e)** Electrophoretic mobility shift assays (EMSA) using labelled JF-549 Rap1-Halo and labelled DNA constructs. **c)** Rap1 binding to DNA containing **P3_*S2***. **d**) Rap1 binding to DNA containing **P3_*S1***. **e**) Rap1 binding to DNA without Rap1 binding site. Titration of Rap1-Halo from 0 – 128 nM and incubated 10mins before loading on native-PAGE. DNA - Rap1-Halo complexes migrate to 1.5 kb and can be seen in both 647 and 549 illumination channels.

**Supplementary Fig. 2.**
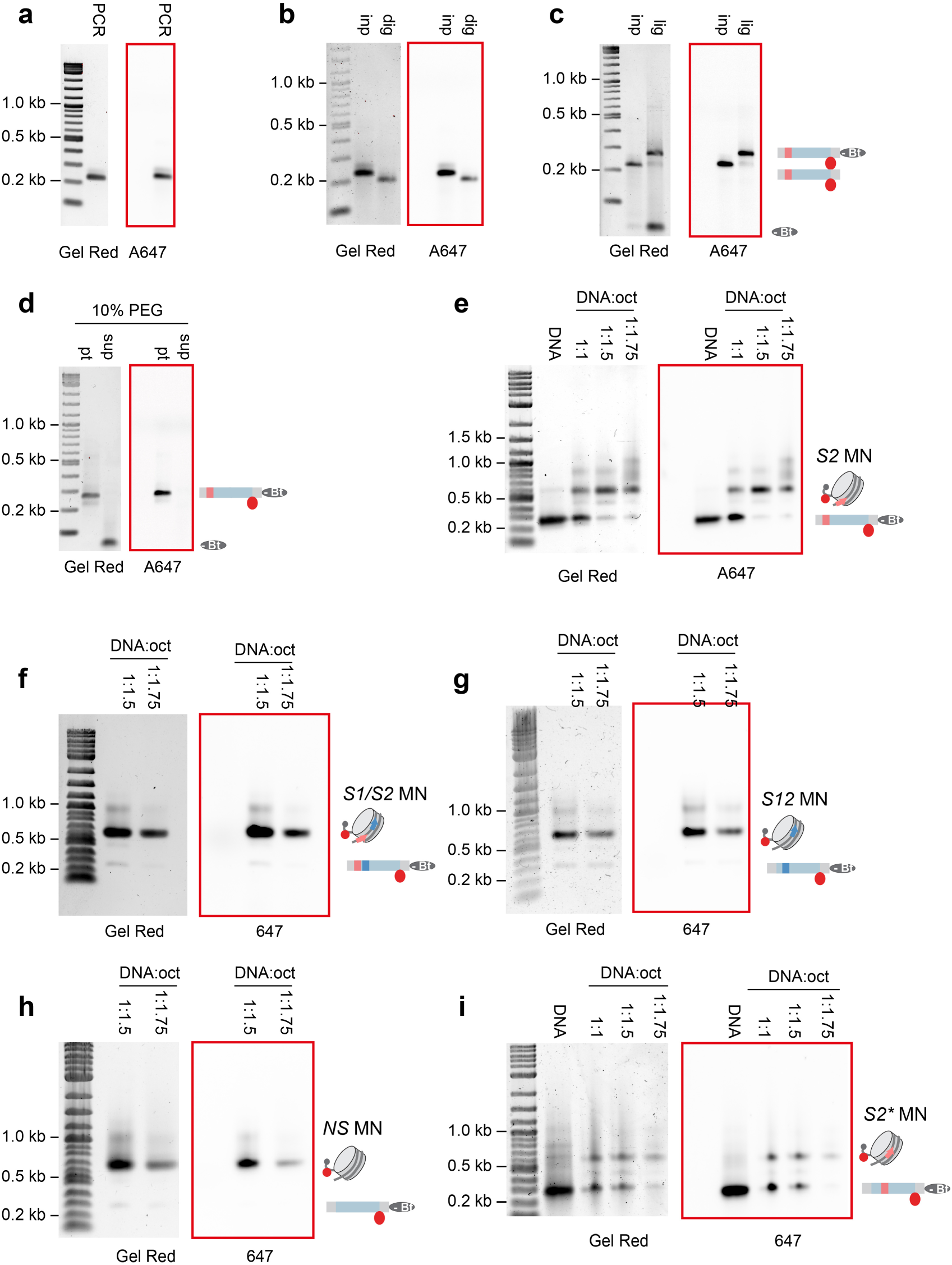
Mononucleosome formation: **a)** Large-scale PCR of labelled **P3_*S2*** DNA. **b)** *DraIII* and *BsaI* digestion of PCR generated **P3_*S2*** DNA. **c)** Ligation of biotin-containing anchor (grey oval) to complementary *BsaI* site on digested PCR construct containing a fluorescent dye (red circle). **d)** PEG purification of biotin ligated DNA construct to remove excess biotin-anchor. **e)** Refolded octamers are titrated into purified **P3_*S2*** DNA and undergo dialysis from 2 M to 0.1 M KCl. These are then run on 0.8% agarose gel in 0.25 TB. Saturation of nucleosomes occurs at DNA to Octamer ratio of 1:1.5. **f)** 0.8% agarose gel in 0.25 TB of nucleosomes containing **P3_*S1S2*** DNA with a biotin anchor. **g)** 0.8% agarose gel in 0.25 TB of nucleosomes containing **P3_*S1*** DNA with a biotin anchor. **h)** 0.8% agarose gel in 0.25 TB of nucleosomes containing **P3** DNA with a biotin anchor. **i)** 0.8% agarose gel in 0.25 TB of nucleosomes containing **P3_*S2*\*** DNA with a biotin anchor.

**Supplementary Fig. 3.**
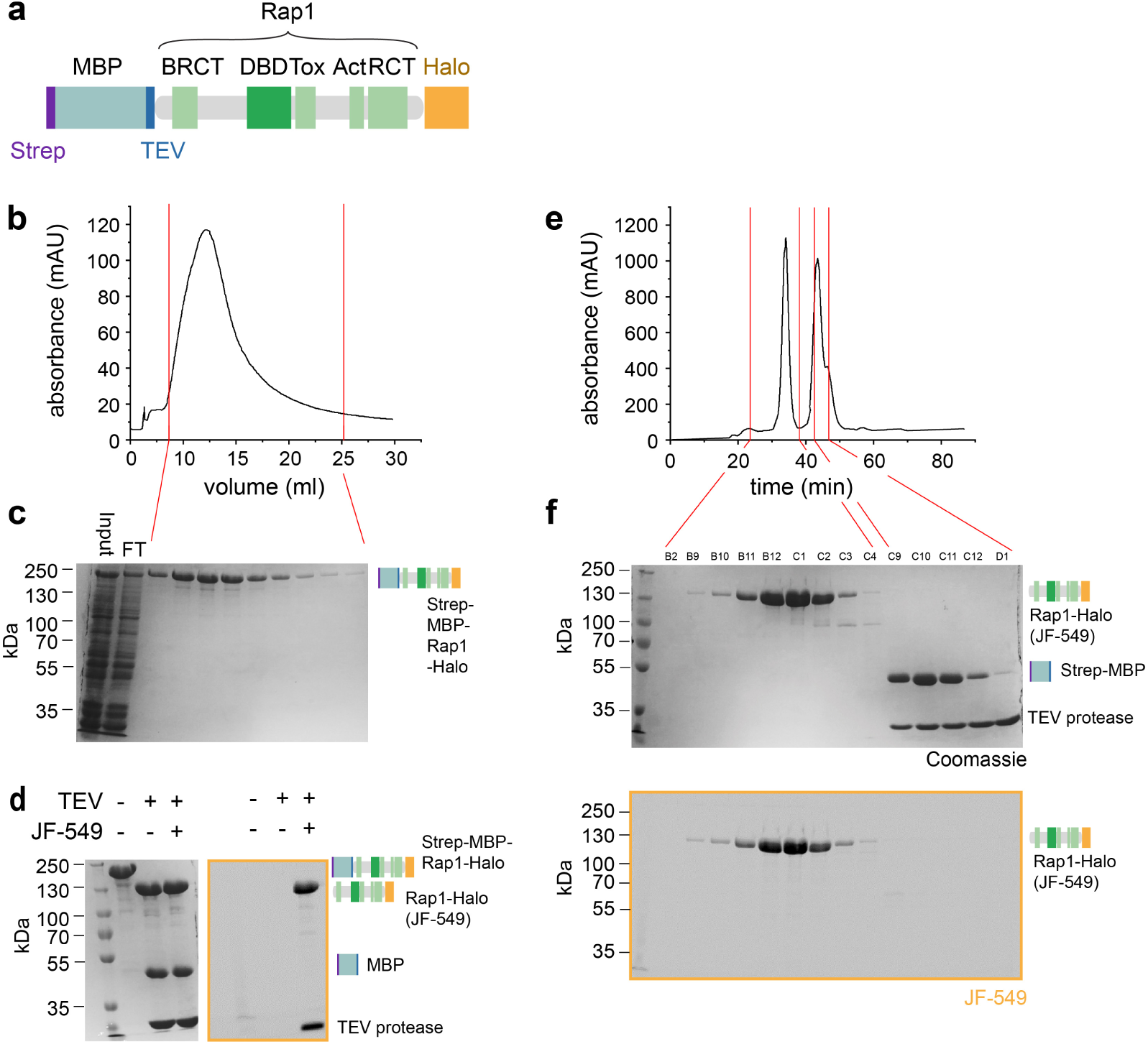
Purification of Rap1: **a)** Schematic representation of full-length baculovirus expressed Rap1-Halo construct. Strep tag (purple), Maltose binding protein (cyan), TEV cleavage site (blue), BRCA 1 C Terminus (BRCT) (green), DNA Binding Domain (DBD) (dark green), Toxicity region (Tox), Transcription Activation domain (Act), Rap1 C-Terminus (RCT) and Halo-Tag (orange). **b-c)** Affinity chromatography profile and SDS-PAGE of corresponding fractions, including supernatant (FT) after centrifugation of lysed cells (input). **d)** SDS-PAGE of TEV protease digestion and Halo-tag labelling using JF-549. **e-f)** Gel filtration profile and SDS-PAGE of corresponding fractions.

**Supplementary Fig. 4.**
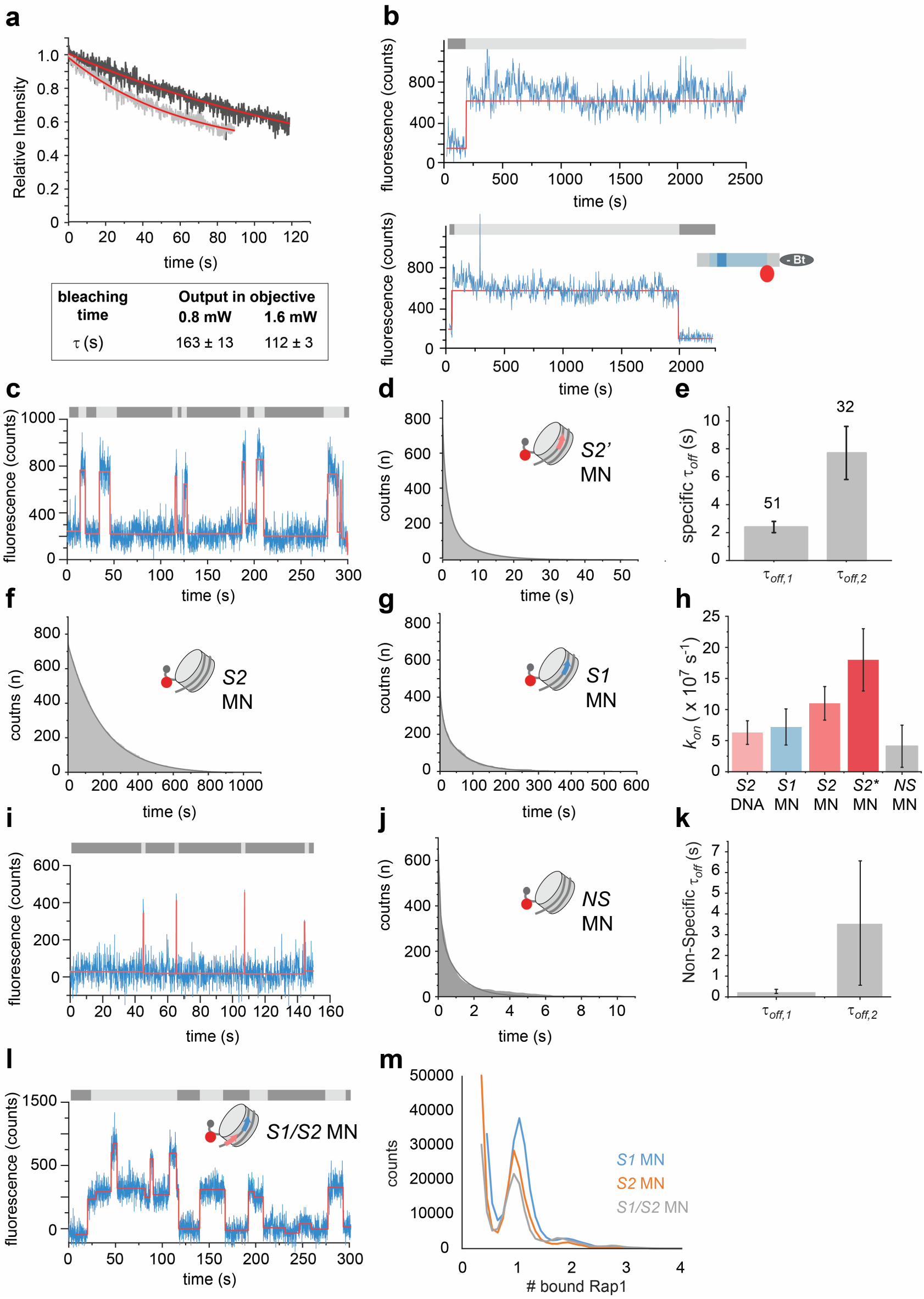
Mononucleosome binding and photobleaching analysis: **a)** Immobilized Rap-JF-549 was continuously illuminated at different power outputs (measured at objective). Intensity decays due to photobleaching were fitted with a single-exponential function yielding the bleaching kinetics of JF-549. **b)** Example traces of biotinylated **P3_*S1*** DNA acquired using smTIRF (blue), with corresponding fit (red). Imagining was done at 0.75 frames/s to limit photobleaching. Individual binding events are seen to last over 30 mins, however a majority of binding events start or finish outside of the measurement window. This reduces the total number of observed events, rendering the acquisition of statistically significant data difficult. **c)** Example traces of biotinylated **P3_*S2*\*** DNA in reconstituted mononucleosomes, acquired using smTIRF (blue), with corresponding fit (red). **d)** Example of a cumulative histogram and tri-exponential fit of mononucleosomes using **P3_*S2*\*** DNA (*S2*\* MN). Data from 100-200 traces **e)** Specific residence times of Rap1 binding to *S2*\* MN. The width of the bar indicates the relative population associated to each time constant. **f)** Example of a cumulative histogram of dark times (*t_dark_*) and mono-exponential fit of mononucleosomes containing **P3_*S2*** DNA (*S2* MN). Data from 200-300 traces. **g)** Example of a cumulative histogram of *t_dark_* and mono-exponential fit of mononucleosomes containing **P3_*S1*** DNA (*S1* MN). Data from 100-200 traces **h)** Association kinetics (*k_on_*) of Rap1 to indicated MNs. **i)** Example trace of biotinylated **P3** 601 DNA (*NS* MN) acquired using smTIRF (blue), with corresponding fit (red). **j)** Example of a cumulative histogram and tri-exponential fit of *NS* MN. Data from 100-200 traces. **k)** Specific residence times of Rap1 binding to *S2* MN. The width of the bar indicates the relative population associated to each time constant. **l**) Example trace of mononucleosomes containing **P3_*S1S2*** (*S1/S2* MN) acquired using smTIRF (blue), with corresponding fit (red), demonstrating a superposition of *S1* and *S2* binding kinetics. **m**) Intensity distribution histograms for mononucleosomes containing either *S1*, *S2* or *S1S2*, reporting on the number of simultaneously bound Rap1 molecules (from 100 traces each). Under the measurement conditions (50 pM Rap1), even when two Rap1 binding sites are present, mostly single-binding events are observed.

**Supplementary Fig. 5.**
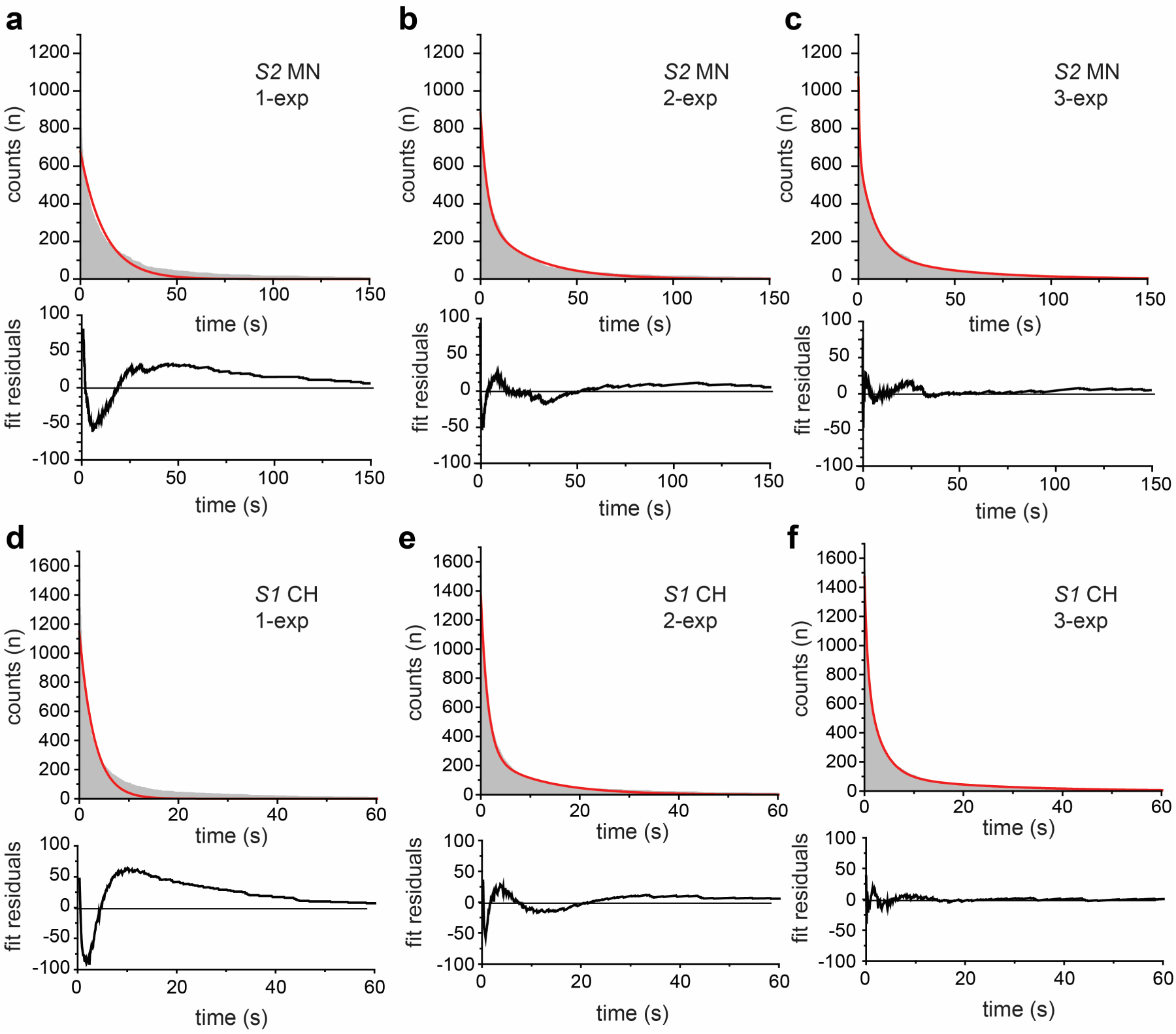
Choice of fitting function: **a)** Example of a cumulative histogram of Rap1 residence times on a **P3_*S2*** mononucleosome (*S2* MN), fit with mono-exponential decay with residuals (below). **b)** Example of a cumulative histogram of Rap1 residence times on a **P3_*S2*** mononucleosome (*S2* MN), fit with bi-exponential decay with residuals (below). **c)** Example of a cumulative histogram of Rap1 residence times on a **P3_*S2*** mononucleosome (*S2* MN), fit with tri-exponential decay with residuals (below). **f)** Example of a cumulative histogram of Rap1 residence times on a **CH_*S1*** chromatin fiber (*S1* CH), fit with mono-exponential decay with residuals (below). **e)** Example of a cumulative histogram of Rap1 residence times on a **CH_*S1*** chromatin fiber (*S1* CH), fit with bi-exponential decay with residuals (below). **f)** Example of a cumulative histogram of Rap1 residence times on a **CH_*S1*** chromatin fiber (*S1* CH), fit with tri-exponential decay with residuals (below).

**Supplementary Fig. 6.**
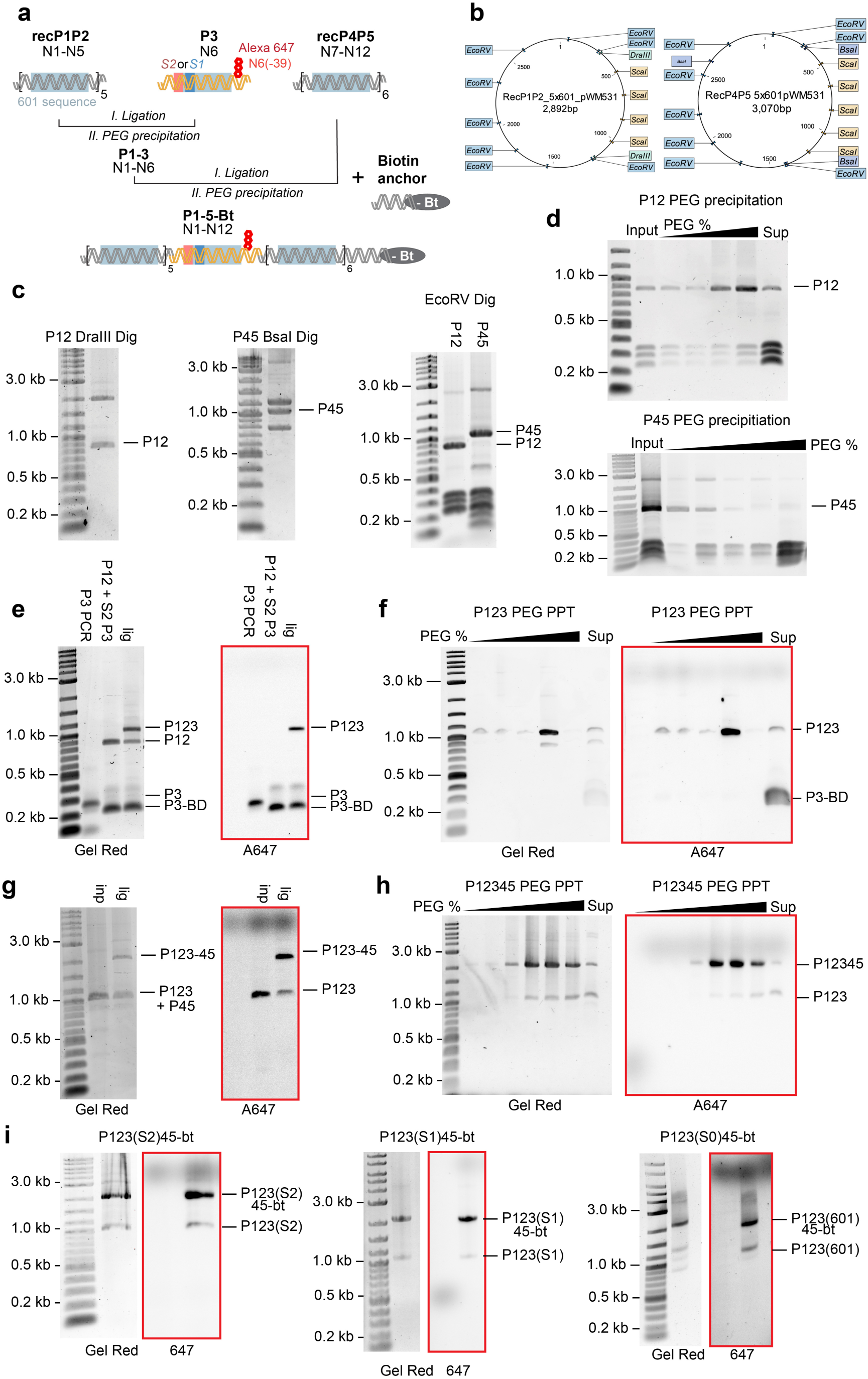
Chromatin assembly: **a)** Schematic representation of the ligation scheme used to generate 12-mer chromatin arrays with 3-piece ligation. Convergent ligation from sequential ligations and PEG purifications of individual pieces. **b)** Scheme of recombinantly expressed rec**P1P2** and rec**P4P5** pieces containing 5×601 and 6×601 sequences respectively. These plasmids are engineered to simplify PEG precipitation purification (*EcoRV*) and nucleosome quality control (*ScaI*). **c)** Example of rec**P1P2** and rec**P4P5** removal from plasmid backbone via successive restriction digestion (*DraIII* and *EcoRV* for rec**P1P2**, *BsaI* and *EcoRV* for rec**P4P5**). **d)** PEG precipitation purification of digested rec**P1P2** and rec**P4P5** pieces by sequentially increasing PEG concentrations. Each iteration consists of centrifugation and removal of supernatant too which PEG is added. **e)** Ligation of individually purified rec**P1P2** and **P3** (**P3_*S1***, **P3_*S2*** and **P3**) pieces, using excess **P3**. **f)** PEG precipitation purification of ligated **P1P2P3** pieces by sequentially increasing PEG concentrations. **g)** Ligation of **P1P2P3** with **P4P5-bt**. **h)** PEG precipitation purification of ligated **P1P2P3P4P5-bt** pieces by sequentially increasing PEG concentrations. **i)** Pooled **P1P2P3P4P5-bt** fractions after PEG purification for **P123_*S2*-45-bt (CH_*S2*)**, **P123_*S1*-45-bt (CH_*S1*)** and **P123(601)45-bt (CH_601)** 12-mer DNA construct.

**Supplementary Fig. 7.**
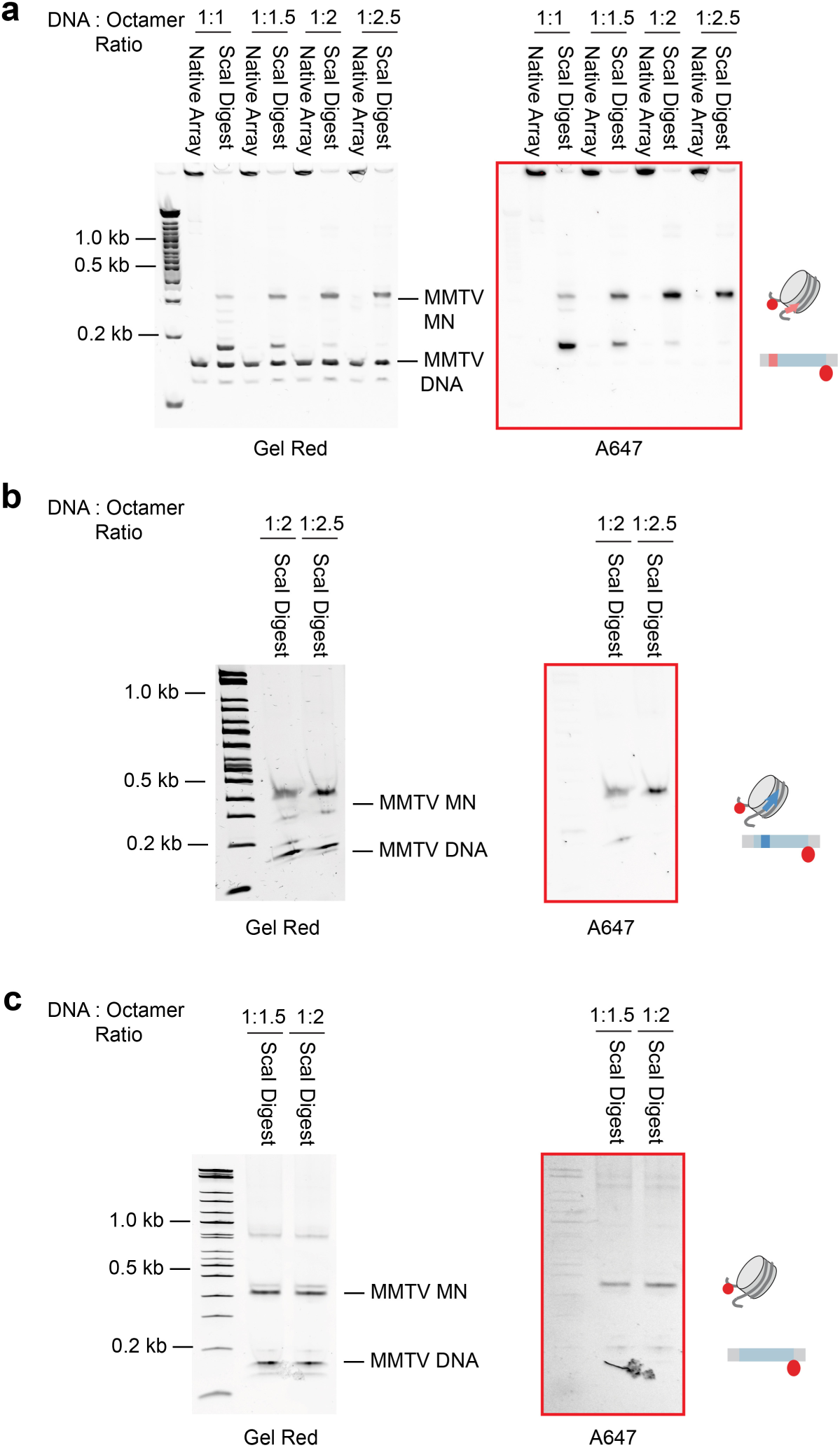
Chromatin assembly: **a)** Analysis of chromatin formation after overnight dialysis using *ScaI* restriction digestion. Different DNA to octamer ratios were tested to find optimal 601 array saturation conditions. MMTV buffer DNA is used as an indicator of saturation of arrays, as 601 saturation precedes the formation of MMTV mononucleosomes. In this gel we see the progressive disappearance of free **CH_*S2*** DNA with the concomitant formation of both 601 and MMTV nucleosomes. We see array saturation at 1:2.5 DNA to octamer ratio. **b)** Gel of **CH_*S1*** Array assembly showing saturation at 1:2.5 DNA to octamer ratio. **c)** Gel of 12-mer **CH_601** Array assembly showing saturation at 1:2 DNA to octamer ratio.

**Supplementary Fig. 8.**
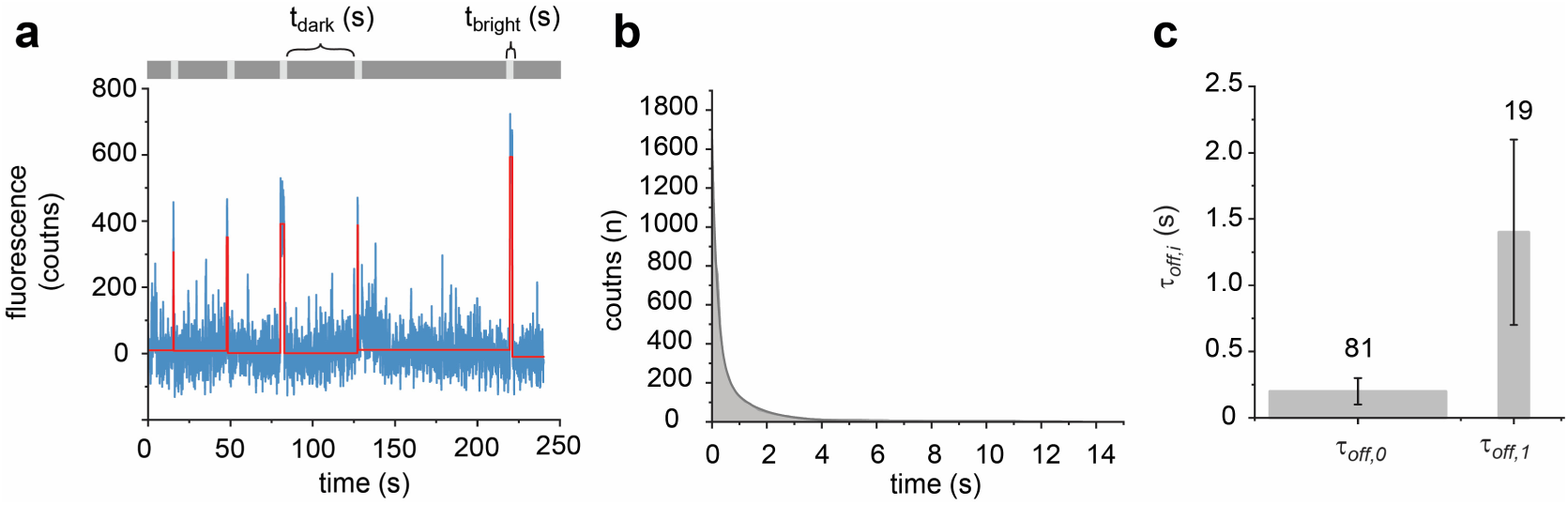
Rap1 binding to CH *NS* chromatin detected by co-localization TIRF. **a)** Example trace from smTIRF experiments using 12-mer **CH *NS*** chromatin without Rap1 site. **b)** Example cumulative histogram of 12-mer **CH *NS*** chromatin smTIRF measurements. The data fit by a bi-exponential decay function. **c)** Residence times for Rap1 on 12-mer **CH *NS*** chromatin showing τ_off, 1_ 0.2 ± 0.1 s and τ_off, 2_ 1.4 ± 0.7 s. The width of the bars indicate the percentage of events associated with the indicated time constants.

**Supplementary Fig. 9.**
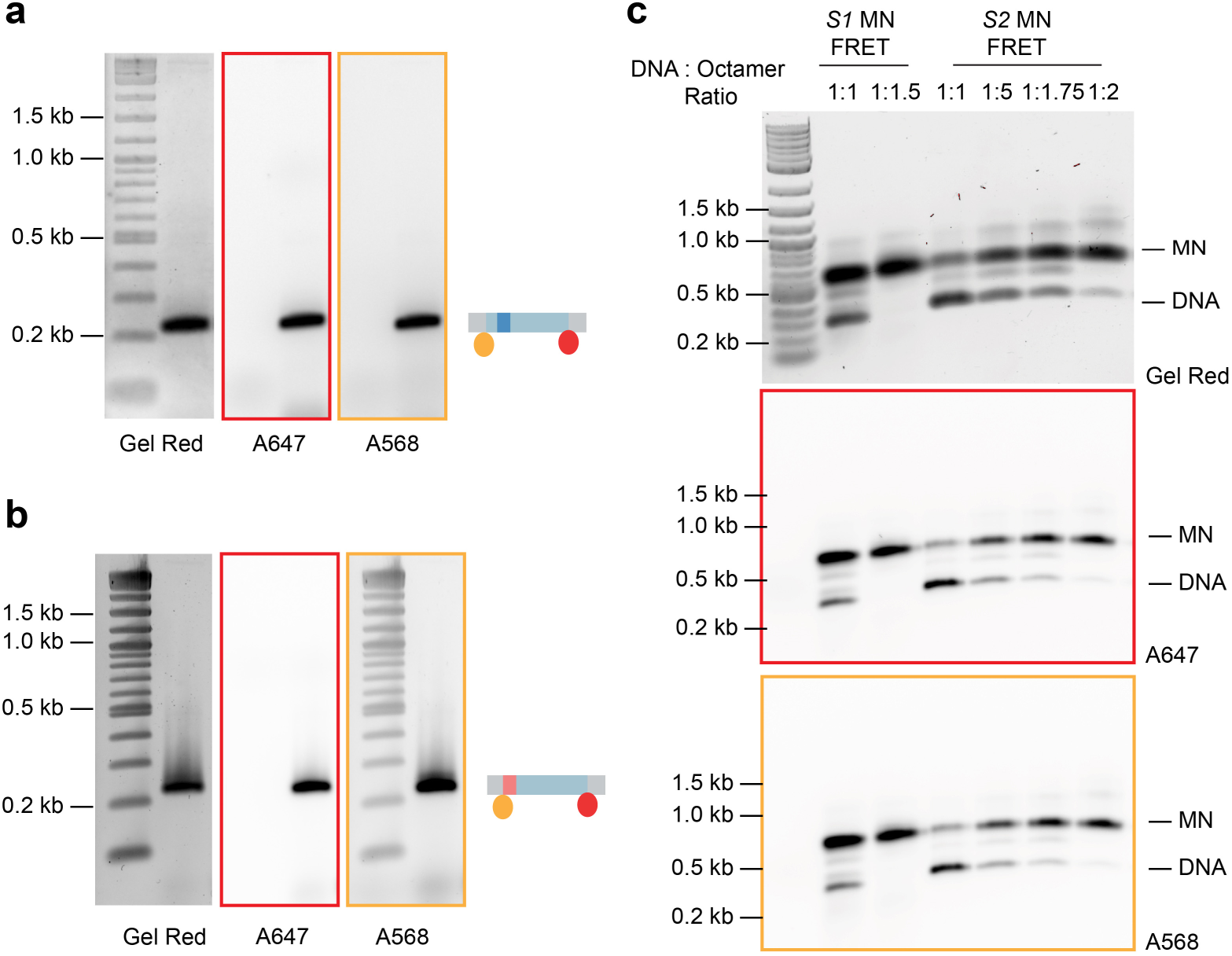
Ensemble FRET nucleosome assembly: **a)** PCR generation of dual labelled 1×601 P3 pieces containing Rap1 *S1* binding site (**MN_*S1*_FRET**). **b)** PCR generation of dual labelled 1×601 P3 pieces containing Rap1 *S2* binding site (**MN_*S2*_FRET**). **c)** Gel showing nucleosome formation by titration of refolded octamers and dialysis. Both *S1* and *S2* nucleosome assembly are visible *S1* shows complete saturation, whereas *S2* nucleosomes contain traces of free DNA.

**Supplementary Fig. 10.**
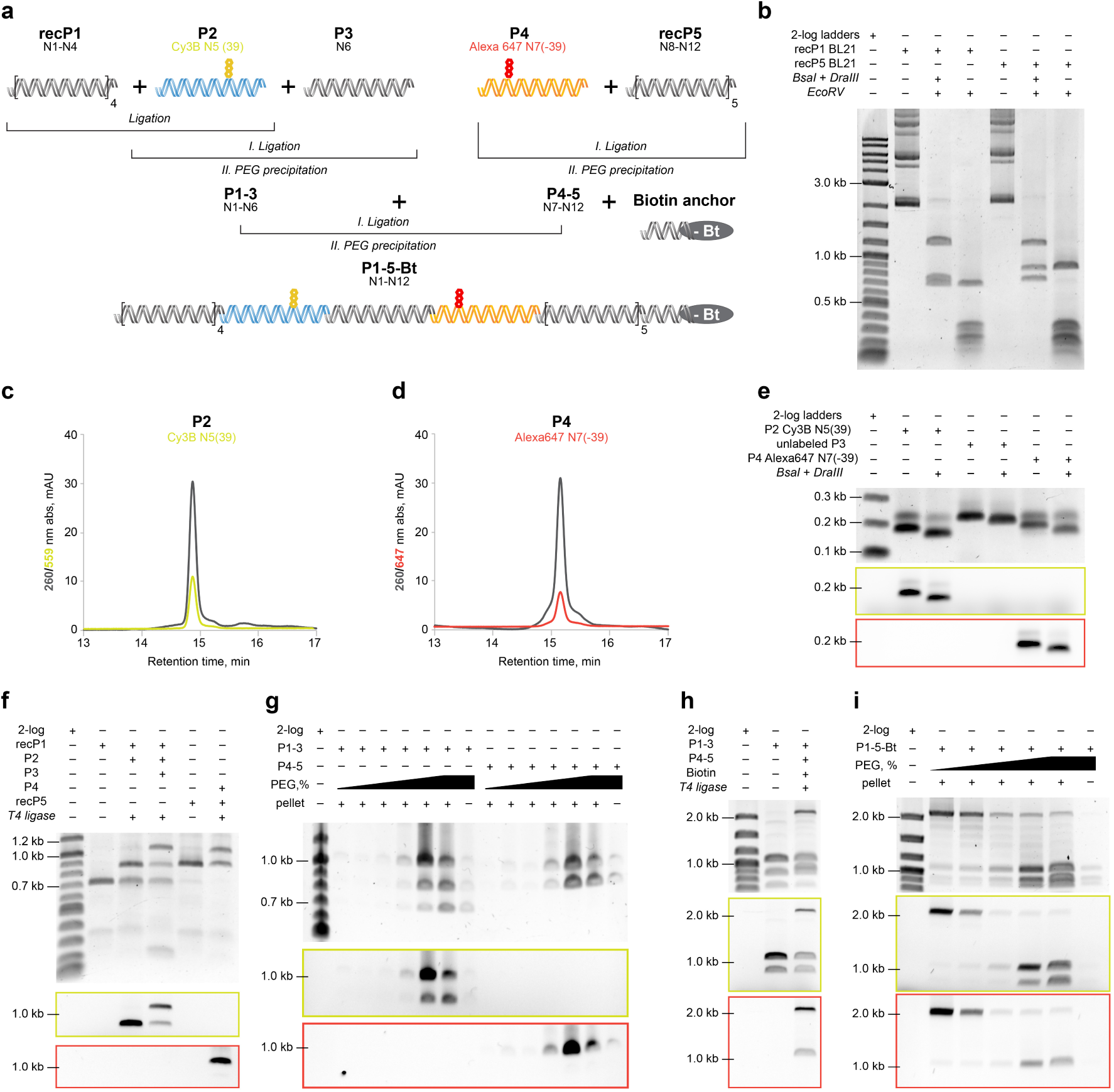
Synthesis of doubly-labeled 12×601 DNA: **a)** Scheme of convergent assembly and purification of doubly-labeled 12×601 DNA from 5 pieces of repeating 601 sequences: recP1 4×601, P2 1×601 labeled by Cy3B (yellow sphere), P3 1×601, P4 1×601 labeled by Alexa Fluor 647 (Alexa647, red sphere), and recP5 5×601. Three first pieces and two latter ones are ligated to form two singly-labeled intermediate 6×601 fragments, P1-3 and P4-5, and ultimately purified from the individual pieces by PEG precipitation. The two intermediate 6×601 pieces and a biotin anchor (-Bt) are ligated to produce the doubly-labeled and biotinylated 12×601 DNA followed by PEG precipitation to separate from the intermediates. **b**) Excision recP1 4×601 (lane 2-4) and recP5 5×601 (lane 5-7) with non-palindromic overhangs by complete digestion first with *BsaI* and *DraIII* followed by plasmid backbone fragmentation by *EcoRV*. **c**, **d**) RP-HPLC profiles of fluorescent labeling on P2 and P4 respectively by Cy3B (yellow) and Alexa647 (red) at the 39-base-pair position relative to the dyad in the 601 sequence. **e)** PCR and digestion, with *BsaI* and *DraIII* to produce unique non-palindromic cohesive ends, of Cy3B-labeled P2 (lane 2-3), unlabeled P3 (lane 4-5) and Alexa647-labeled P4 (lane 6-7). **f**, **g**) Ligation and PEG purification of the singly-labeled intermediate 6×601 fragments: P1-3 (lane 2-4**f** and lane 2-8**g**), and P4-5 (lane 5-6**f** and lane 9-15**g**). The enriched intermediates, P1-3 lane 6-7 and P4-5 lane 12-14, were collected for the final ligation. **h**, **i**) Final ligation and PEG purification of the doubly-labeled biotinylated 12×601 DNA. The enriched final products (lane 2-3**i**) were collected. All agarose gels were imaged using fluorescence imaging of Cy3 (yellow frame) and Alexa647 (red frame), stained and imaged with GelRed (no border frame).

**Supplementary Fig. 11.**
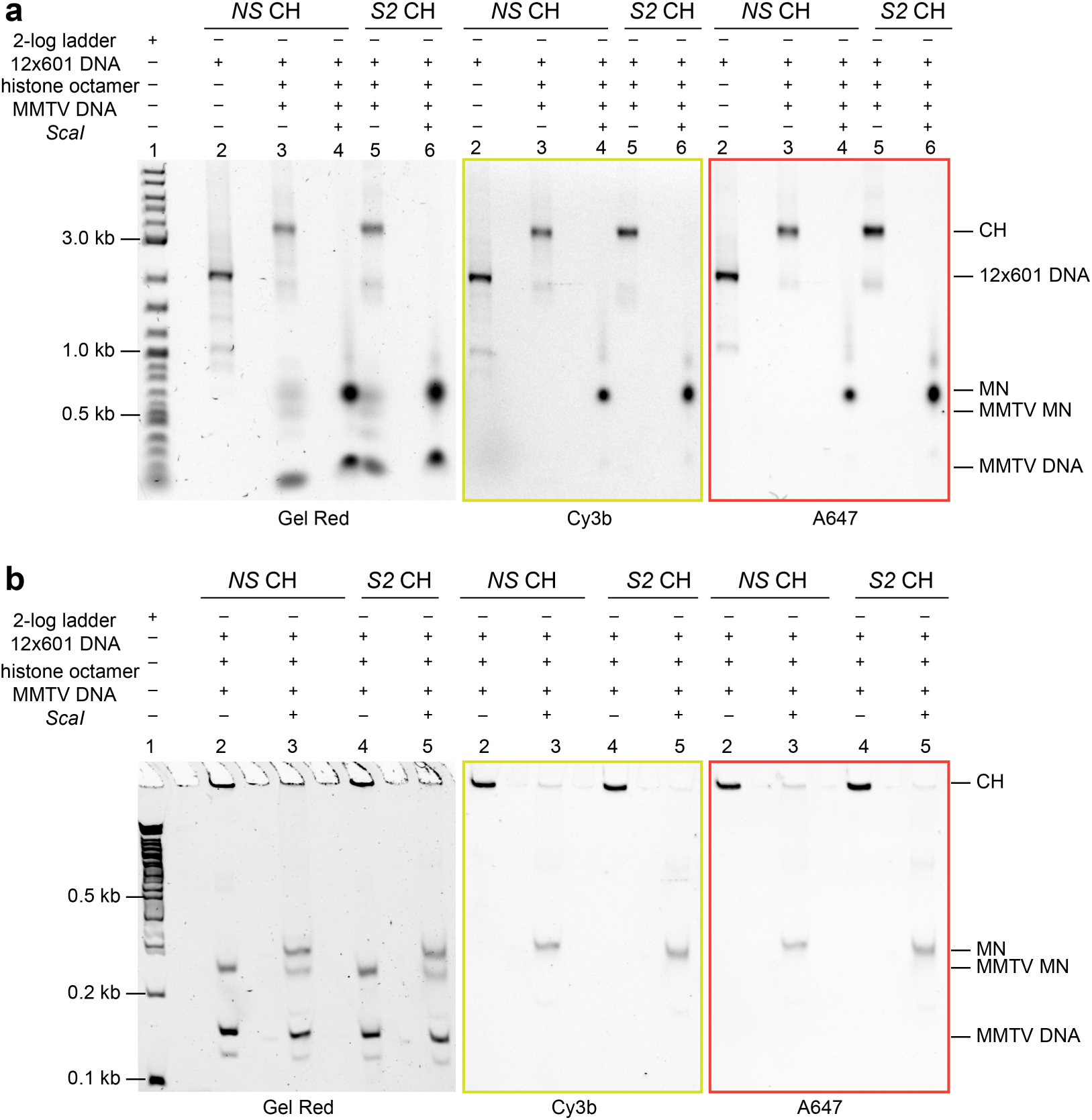
Biochemical analysis of chromatin assembly by gel electrophoresis: **a,b**) Agarose gel and 5% TBE polyacrylamide analysis of chromatin assembled from doubly-labeled biotinylated 12×601 DNA (**CH_*S2*_FRET and CH_601_FRET**) and wild-type histone octamer. To avoid overloading array DNA with histone octamers, low-affinity buffer DNA, MMTV DNA, was added, resulting in the formation of a small amount of buffer nucleosomes, MMTV nucleosomes, as a signature of saturated chromatin fibers (lanes 3&5**a**, and lanes 2&4**b**). Digestion of chromatin fibers with the restriction enzyme *ScaI* liberates mononucleosomes (lanes 4&6**a**, and lanes 3&5**b**). All gels were imaged using fluorescence imaging of Cy3 (yellow frame) and Alexa647 (red frame), stained and imaged with GelRed (no border frame).

**Supplementary Fig. 12.**
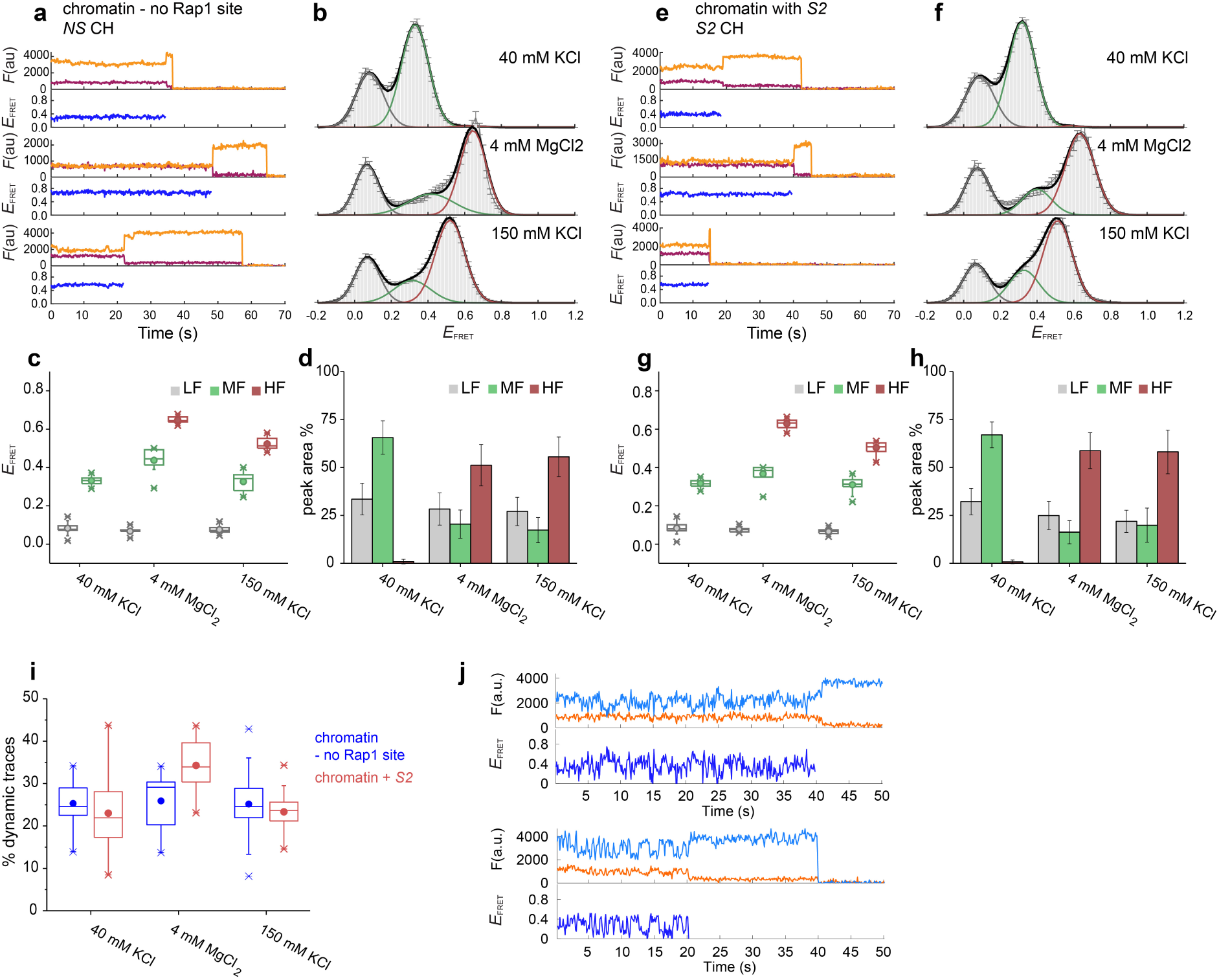
Characterization of chromatin compaction by smFRET: **a**, **e**) Individual traces of donor (green), acceptor (red) and FRET efficiency (*E_FRET_*), and (**b**, **f**) histograms of *E_FRET_* of a single chromatin fibers in 40 mM KCl, 4 mM MgCl_2_ and 150 mM KCl as indicated, error bars shown as s.e.m, number of traces and parameters of Gaussian fits shown in **Supplementary Table 4. c**, **g**) Peak center and (**d**, **h**) percentage of the integral area of Gaussian fits with LF (grey, peak *E_FRET_* < 0.2), MF (green, 0.2 ≤ *E_FRET_*≤ 0.4) and HF (red, *E_FRET_* > 0.4). Experiments were performed on control chromatin (*NS* CH, **a**-**d**) in parallel with chromatin containing Rap1 site 2 (*S2* CH, **e**-**h**). **i**) Percentage of dynamic traces seen in two types of chromatin. **j**) Examples of dynamic traces, showing anticorrelated donor and acceptor fluorescence fluctuations, indicating conformational dynamics. Data are shown for chromatin fibers with *S2* (*S2* CH) in the presence of 200 pM Rap1.

**Supplementary Fig. 13.**
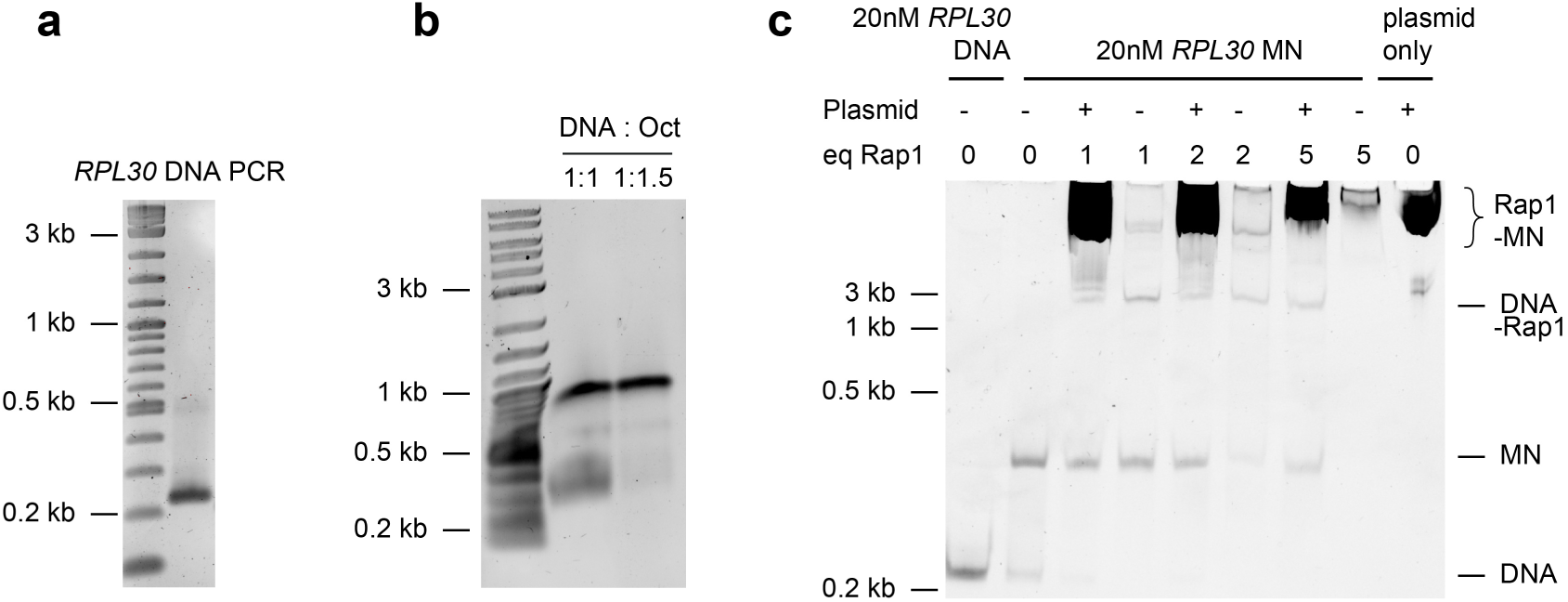
*RPL30* Nucleosome stability: **a)** Large-scale PCR of labelled **P3_*RPL30*** DNA. **b)** Refolded octamers are titrated into purified **P3_*RPL30*** DNA and undergo dialysis from 2 M to 0.1 M KCl. These are then run on 0.8% agarose gel in 0.25 TB. Saturation of nucleosomes occurs at DNA to Octamer ratio of 1:1.5. **c)** To assay *RPL30* nucleosome stability, Rap1 was titrated to 1, 2 and 5 equivalents of *RPL30* mononucleosome and left at room temperature for 10 mins. To recover the nucleosomes (that were bound by Rap1), buffer DNA was added. Nucleosome integrity is ensured as no sub-nucleosomal bands (below the nucleosome band, NCP) are observed. A minor amount of free DNA forms Rap1-DNA adducts.

**Supplementary Fig. 14.**
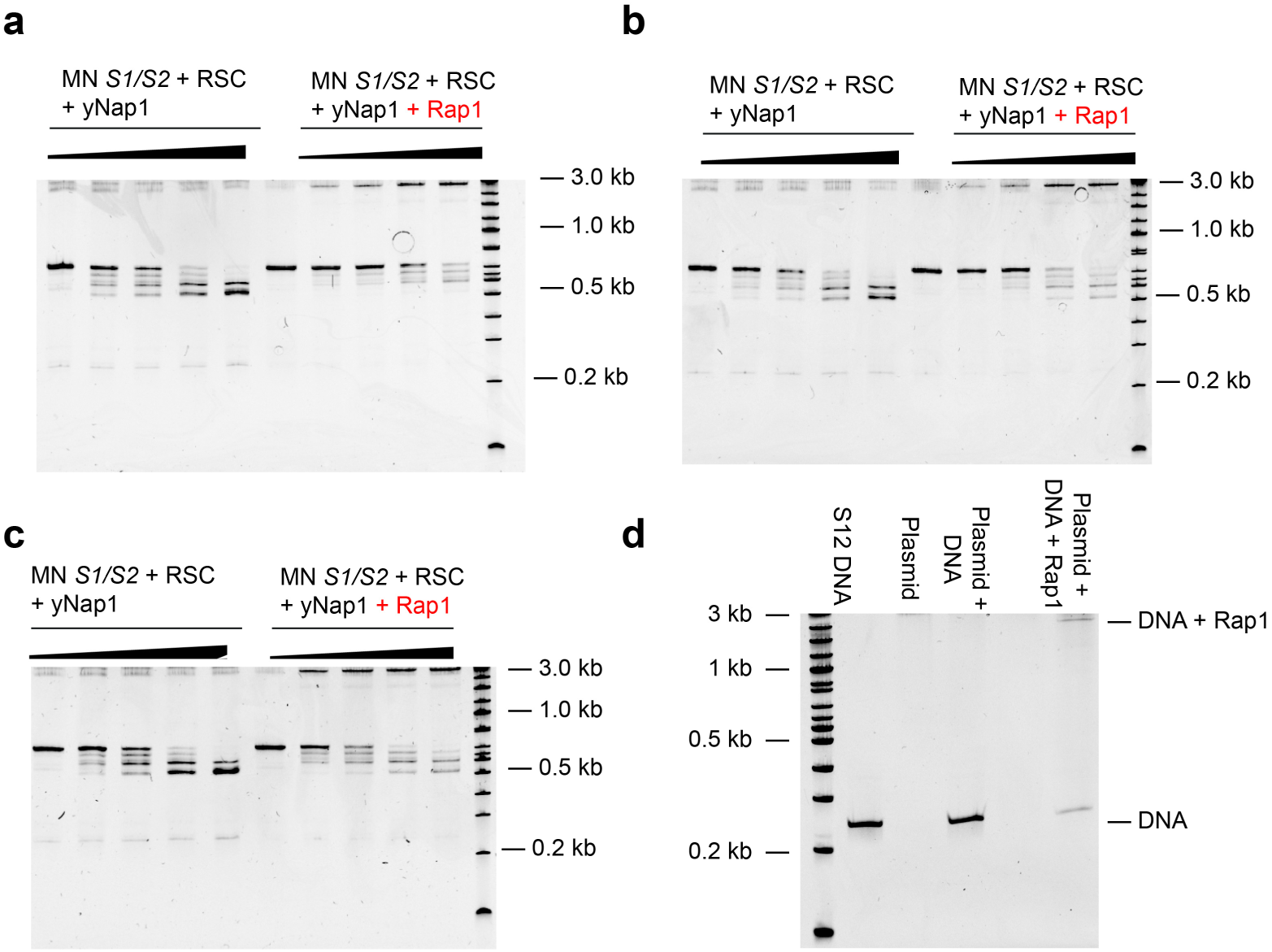
RSC remodeling in the presence of Rap1. **a-c)** Replicates of gels to show RSC mononucleosome remodeling in the presence and absence of Rap1. **d)** Gel showing the identification of the upper-band (Rap1 + DNA), this experiment was done using free *S1*2 DNA (**P3_*S1S2***) and not *S1*2 nucleosomes.

**Supplementary Fig 15.**
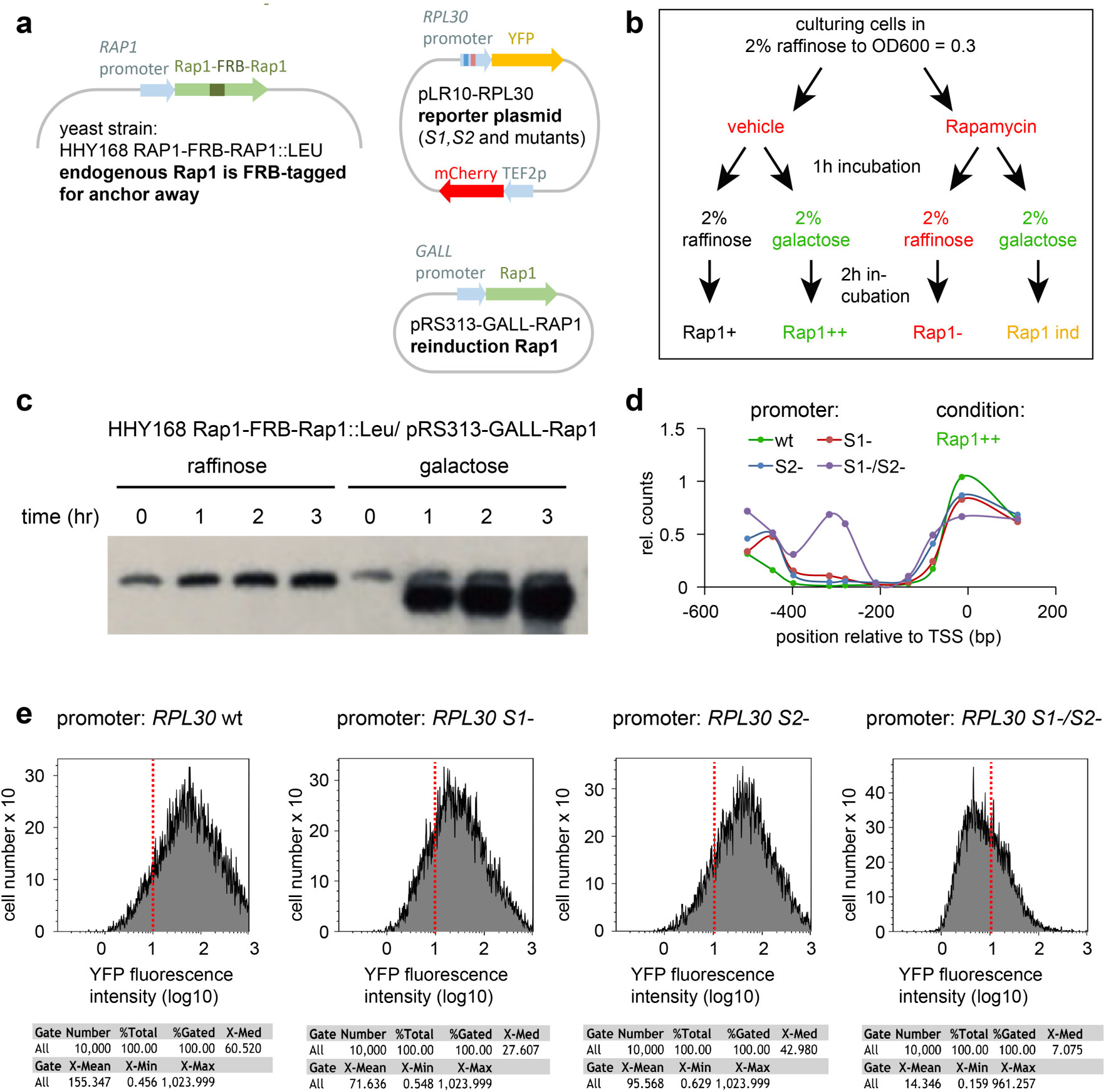
Rap1 chromatin remodeling dynamics in yeast cells. **a**) Yeast constructs: The yeast strain used expresses Rap1 tagged with FRB, which allows to anchor away Rap1 (deplete from the nucleus) upon addition of rapamycin (as anchor serves the FKBP12-tagged plasma membrane protein Pma1p). In the absence of rapamycin, Rap1 binds to the *RPL30* promoter in the reporter plasmid (potentially containing mutated sites *S1* or *S2*, for sequences see **Supplementary Table 6**). Rap1 then induces the reporter gene YFP (Rap1+). Rapamycin addition results in removal of Rap1 (Rap1-). Growth of the yeast cells on galactose results in reinduction of Rap1 from a plasmid under a galactose sensitive promoter (Rap1 ind). **b**) Experimental scheme: Sequential treatment with rapamycin vs. vehicle and galactose vs. raffinose results in 4 samples that are further analyzed by MNase nucleosome mapping and YFP expression by FACS. **c**) Rap1 expression levels before and after induction with galactose. **d**) MNase mapping of different *RPL30* promoter mutants in the Rap1++ sample (for other samples, see **Fig. 6d**). **e**) Determination of reporter gene induction by FACS sorting for YFP fluorescence. Promoter types wt and *S2*-show high YFP expression, for *S1*-expression is reduced and for *S1-/S2-* expression is strongly suppressed.

**Supplementary Table 1.**
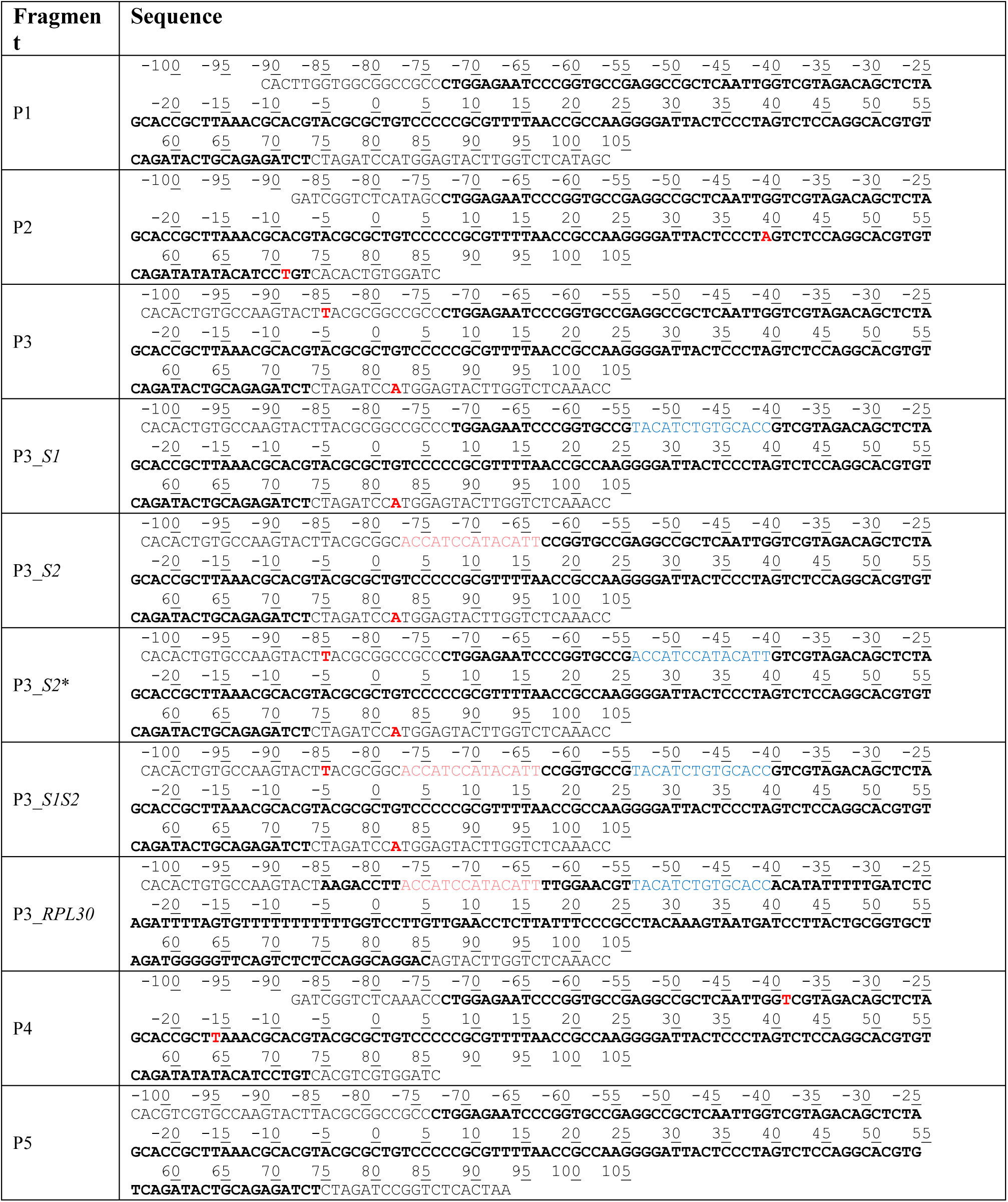
Sequences of 1 × 601 pieces for recombinant and PCR-generated pieces. 601 or native *RPL30* sequences indicated in bold. The labeled base pairs are indicated in red. The numbering is given as number of base-pairs relative to the dyad in the 601 sequence. Rap1 binding sites are indicated in blue for *S1* and burgundy for *S2*.

**Supplementary Table 2.**
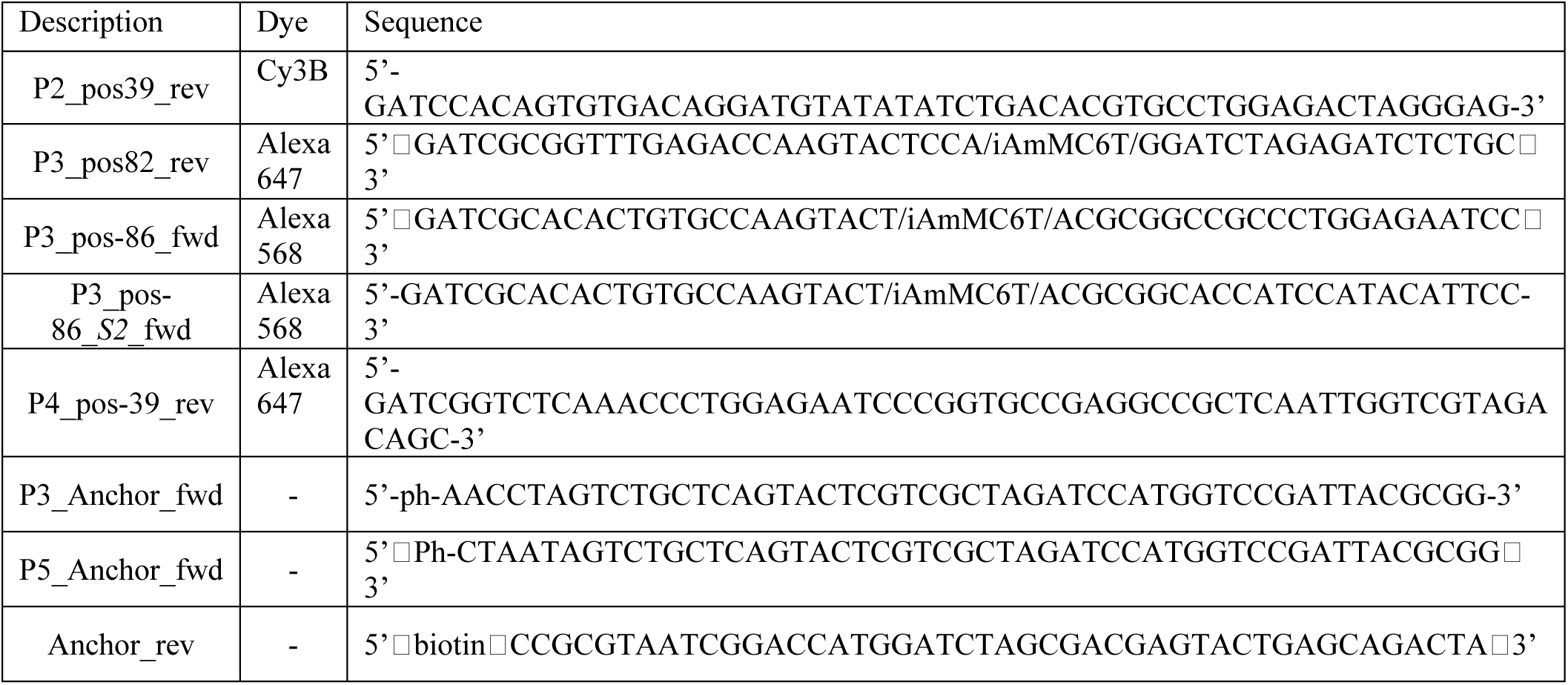
Sequences of all labeled oligonucleotides.

**Supplementary Table 3.** Overview of all chromatin DNA with different combinations of labels.

| Experiment | Name | Backbone | Dye | Modification |
| --- | --- | --- | --- | --- |
| Rap1 nucleosome binding | MN S1 | P3_S1 | Alexa647 (82) | 3' biotin |
|  | MN S2 | P3_S2 | Alexa647 (82) | 3' biotin |
|  | MN S2* | P3_S2* | Alexa647 (82) | 3' biotin |
|  | MN S1/S2 | P3_S1S2 | Alexa647 (82) | 3' biotin |
|  | MN NS | P3 | Alexa647 (82) | 3' biotin |
| Rap1 chromatin binding | CH S1 | P1P2 ; P3_S1 ; P4P5 | Alexa647 (82) | 3' biotin |
|  | CH S2 | P1P2 ; P3_S2 ; P4P5 | Alexa647 (82) | 3' biotin |
|  | CH NS | P1P2 ; P3 ; P4P5 | Alexa647 (82) | 3' biotin |
| Nucleosome FRET | MN S1 FRET | P3_S1 | P3: Alexa647 (82), Alexa568 (-86) | 3' biotin |
|  | MN S2 FRET | P3_S2 | P3: Alexa647 (82), Alexa568 (-86) | 3' biotin |
| Chromatin FRET | CH NS FRET | P1 ; P2 ; P3 ; P4 ; P5 | P2: Cy3B (39), P4: Alexa647 (-39) | 3' biotin |
|  | CH S2 FRET | P1 ; P2 ; P3_S2 ; P4 ; P5 | P2: Cy3B (39), P4: Alexa647 (-39) | 3' biotin |
| Ensemble binding & RSC experiments | MN RPL30 | P3_RPL30 |  |  |
|  | MN S1/S2 | P3_S1S2 |  |  |

**Supplementary Table 4.** Summary of Gaussian fits and percentage of dynamic traces from chromatin compaction experiments.

|  |  | CH NS FRET |  |  | CH S2 FRET |  |  |
| --- | --- | --- | --- | --- | --- | --- | --- |
|  |  | 40 mM KCl | 4 mM Mg<sup>2+</sup> | 150 mM KCl | 40 mM KCl | 4 mM Mg<sup>2+</sup> | 150 mM KCl |
| LF | A<sub>1</sub> | 0.04 ± 0.01 | 0.03 ± 0.01 | 0.03 ± 0.01 | 0.04 ± 0.01 | 0.03 ± 0.01 | 0.03 ± 0.01 |
|  | c<sub>1</sub> | 0.08 ± 0.03 | 0.07 ± 0.02 | 0.08 ± 0.02 | 0.08 ± 0.03 | 0.08 ± 0.01 | 0.07 ± 0.01 |
|  | σ<sub>1</sub> | 0.07 ± 0.01 | 0.07 ± 0.01 | 0.07 ± 0.01 | 0.07 ± 0.01 | 0.06 ± 0.01 | 0.07 ± 0.01 |
|  | % area | 33.5 ± 8.3 | 28.3 ± 8.4 | 27.1 ± 7.4 | 32.2 ± 6.9 | 24.9 ± 7.4 | 22 ± 5.8 |
| MF | A<sub>2</sub> | 0.07 ± 0.01 | 0.02 ± 0.01 | 0.02 ± 0.01 | 0.07 ± 0.01 | 0.02 ± 0.01 | 0.02 ± 0.01 |
|  | c<sub>2</sub> | 0.33 ± 0.02 | 0.44 ± 0.06 | 0.33 ± 0.05 | 0.32 ± 0.02 | 0.37 ± 0.05 | 0.31 ± 0.04 |
|  | σ<sub>2</sub> | 0.07 ± 0.01 | 0.07 ± 0.01 | 0.08 ± 0.01 | 0.07 ± 0.01 | 0.08 ± 0.01 | 0.07 ± 0.01 |
|  | % area | 65.6 ± 8.8 | 20.5 ± 7.3 | 17.4 ± 6.6 | 67 ± 6.7 | 16.3 ± 6 | 19.9 ± 8.9 |
| HF | A<sub>3</sub> | 0 | 0.07 ± 0.02 | 0.06 ± 0.01 | 0 | 0.06 ± 0.02 | 0.06 ± 0.01 |
|  | c<sub>3</sub> | 0.55 ± 0.06 | 0.65 ± 0.02 | 0.52 ± 0.03 | 0.58 ± 0.13 | 0.63 ± 0.03 | 0.5 ± 0.03 |
|  | σ<sub>3</sub> | 0.03 ± 0.03 | 0.06 ± 0.01 | 0.07 ± 0.01 | 0.03 ± 0.02 | 0.08 ± 0.02 | 0.08 ± 0.01 |
|  | % area | 0.9 ± 1.2 | 51.2 ± 10.8 | 55.6 ± 10.3 | 0.8 ± 1 | 58.8 ± 9.3 | 58.1 ± 11.4 |
| N total traces |  | 1189 | 595 | 1477 | 1409 | 521 | 1523 |
| N dynamic traces |  | 306 | 154 | 375 | 332 | 175 | 355 |
| % dynamic traces |  | 25.3 ± 5.6 | 25.9 ± 6.9 | 25.2 ± 7.8 | 23 ± 9.1 | 34.3 ± 6 | 23.3 ± 4.3 |
| N of repeats |  | 21 | 10 | 23 | 24 | 10 | 22 |

**Supplementary Table 5.**
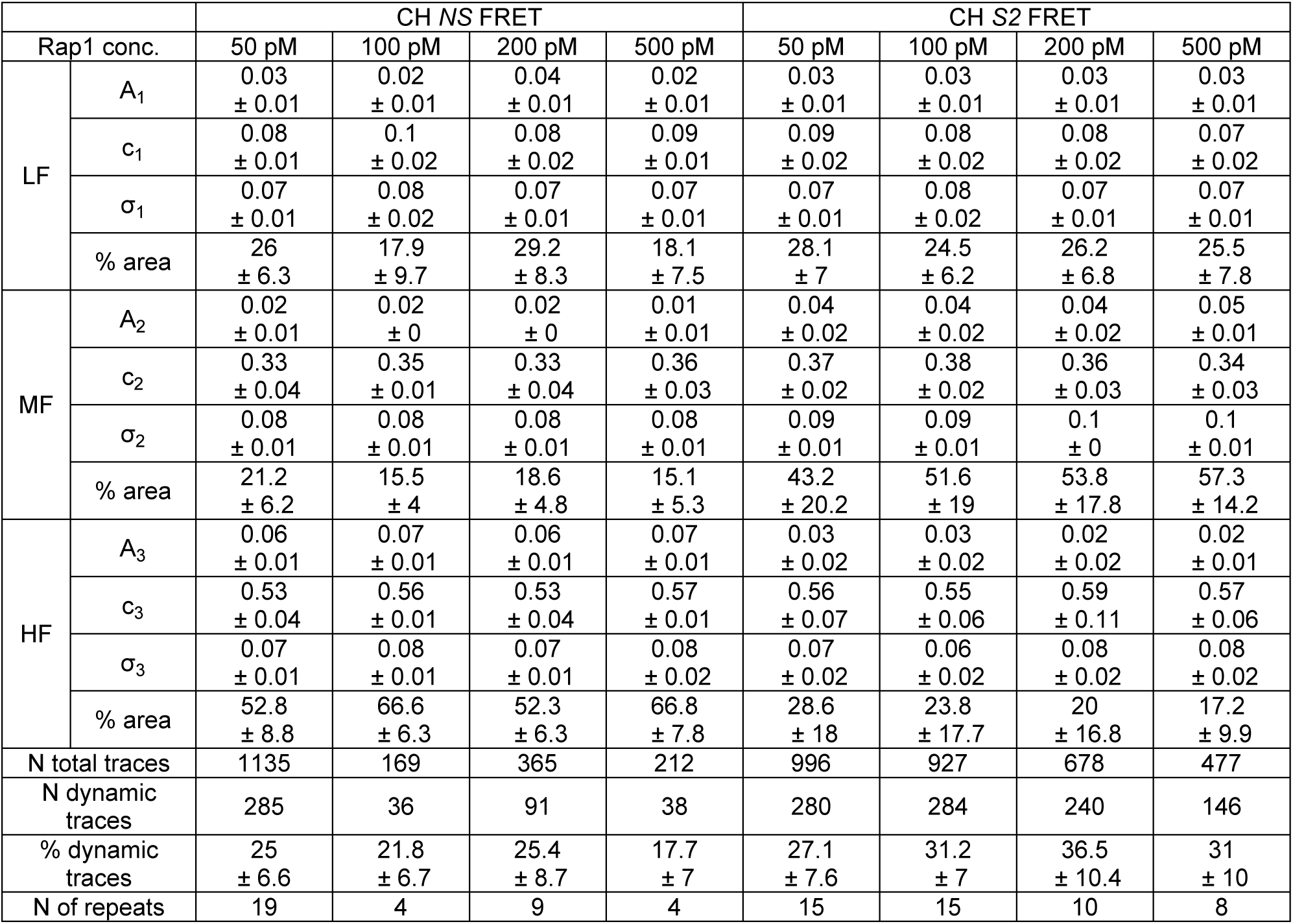
Summary of Gaussian fits and percentage of dynamic traces from chromatin remodeling induced by Rap1 invasion experiments.

**Supplementary Table 6.**
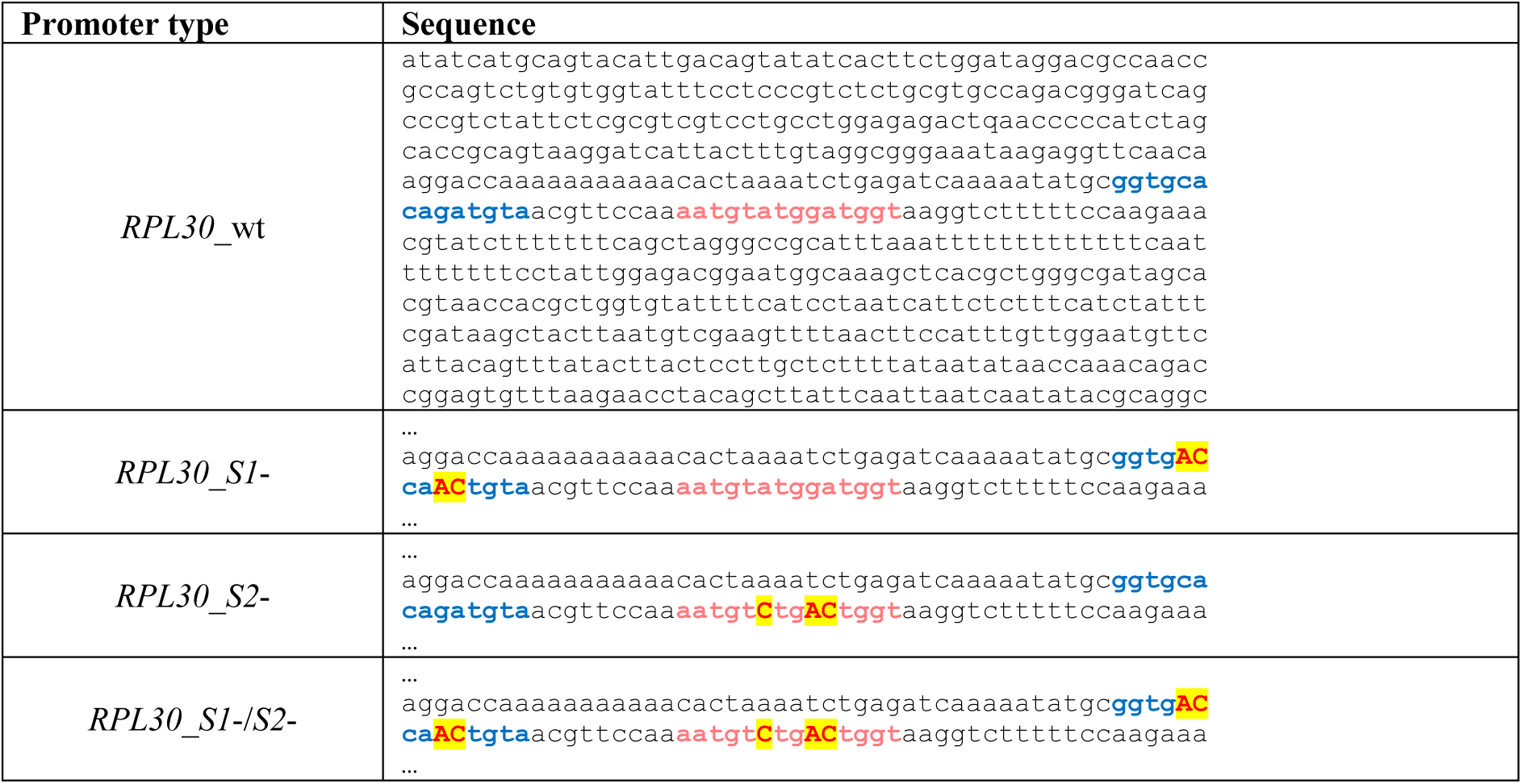
*RPL30* promoter sequences in reporter plasmid for yeast experiments. Rap1 binding site *S1* is indicated in blue, *S2* in burgundy. Mutated residues are given in red and highlighted in yellow.

